# Sex-specific mechanisms of cerebral microvascular BK_Ca_ dysfunction in a mouse model of Alzheimer’s disease

**DOI:** 10.1101/2023.06.06.543962

**Authors:** Josiane F. Silva, Felipe D. Polk, Paige E. Martin, Stephenie H. Thai, Andrea Savu, Matthew Gonzales, Allison M. Kath, Michael T. Gee, Paulo W. Pires

## Abstract

**Background:** Cerebral microvascular dysfunction and nitro-oxidative stress are present in patients with Alzheimer’s disease (AD) and may contribute to disease progression and severity. Large conductance Ca^2+^-activated K^+^ channels (BK_Ca_) play an essential role in vasodilatory responses and maintenance of myogenic tone in resistance arteries. BK_Ca_ impairment can lead to microvascular dysfunction and hemodynamic deficits in the brain. We hypothesized that reduced BK_Ca_ function in cerebral arteries mediates microvascular and neurovascular responses in the *5x-FAD* model of AD.

**Methods:** BK_Ca_ activity in the cerebral microcirculation was assessed by patch clamp electrophysiology and pressure myography, in situ Ca^2+^ sparks by spinning disk confocal microscopy, hemodynamics by laser speckle contrast imaging. Molecular and biochemical analyses were conducted by affinity-purification assays, qPCR, Western blots and immunofluorescence.

**Results:** We observed that pial arteries from 5-6 months-old male and female *5x-FAD* mice exhibited a hyper-contractile phenotype than wild-type (WT) littermates, which was linked to lower vascular BK_Ca_ activity and reduced open probability. In males, BK_Ca_ dysfunction is likely a consequence of an observed lower expression of the pore-forming subunit BK_α_ and blunted frequency of Ca^2+^ sparks, which are required for BK_Ca_ activity. However, in females, impaired BK_Ca_ function is, in part, a consequence of reversible nitro-oxidative changes in the BK_α_ subunit, which reduces its open probability and regulation of vascular tone. We further show that BK_Ca_ function is involved in neurovascular coupling in mice, and its dysfunction is linked to neurovascular dysfunction in the model.

**Conclusion:** These data highlight the central role played by BK_Ca_ in cerebral microvascular and neurovascular regulation, as well as sex-dependent mechanisms underlying its dysfunction in a mouse model of AD.

## Introduction

Alzheimer’s disease (AD) is a neurodegenerative disorder associated with global cognitive decline that affects more than 6 million people in the United States alone^1,2^, and approximately 2/3 of all AD cases are women^1,3^. Alarmingly, it is projected that more than 130 million people will live with AD worldwide by 2050^1,3^. One of the hallmarks of AD is cerebral amyloid angiopathy (CAA), characterized by perivascular accumulation of amyloid-β, which is present in more than 80% of all AD patients^4^. This accumulation is proposed to induce microvascular fragility, contributing to the development of intracerebral hemorrhages^5^, loss of blood-brain-barrier integrity ^6^, and blood flow deficits^7^. These deficits may be a consequence of microvascular dysfunction, which impairs cerebrovascular reserve and neurovascular coupling^8^, a vital process that matches local hyperemia to increased neuronal activity^9^. The ability of the cerebral microcirculation to accommodate these second-to-second changes in local perfusion depends on two main phenomena: the extent of myogenic tone (the intrinsic contractile property of vascular smooth muscle cells that stipulates the vasodilatory reserve) and mechanisms underlying vasorelaxation. Both mechanisms are dependent on the membrane potential of smooth muscle cells: depolarization favors constriction, whereas hyperpolarization favors dilation. Thus, regulation of ion flux is paramount for microvascular function, particularly modulation of K^+^ permeability, of which the large-conductance Ca^2+^-activated K^+^ channel (BK_Ca_) is central.

BK_Ca_ are widely expressed channels that play a critical role in regulating arterial tone by conducting hyperpolarizing currents in vascular smooth cells to oppose vasoconstriction^10,11^. Opening of BK_Ca_ channels occurs downstream from localized intracellular Ca^2+^ release from ryanodine receptors located in the sarcoplasmic reticulum (Ca^2+^ sparks), which activates BK_Ca_ channels, leading to hyperpolarization and dilation^10,12^. BK_Ca_ channels are composed of two subunits, a selective pore (BK_α_) and regulatory (BK_β1,_ BK_γ_) subunits^13^. Further, BK_Ca_ undergoes extensive pre- and post-translational modifications (PTM), including phosphorylation^14–16^, oxidation and nitrosylation, all affecting channel open probability (Po) and conductance^16^. In some tissues, oxidation increases channel activity^17,18^, whereas, in others, it markedly inhibits BK_Ca_^19–21^. Similarly, nitrosylation of thiol moieties due to an electrophilic attack by peroxynitrite, formed by a reaction between nitric oxide (NO) and superoxide (O_2_^−^) in a pro-oxidative environment^22–24^, as seen in AD^25–27^, inhibits BK_Ca_ currents^19^. Despite recent evidence of BK_Ca_ impairment in male APP23 mice^28^, it remains unknown if the reduction in cerebral microvascular BK_Ca_ occurs in females *via* similar mechanisms, and whether neurovascular dysfunction accompanies loss of BK_Ca_ activity. Using a combination of molecular and biochemical tools, ex vivo patch clamp electrophysiology and microvascular pressure myography, and *in vivo* hemodynamic assessments, we show that BK_Ca_ dysfunction mediates cerebral microvascular and neurovascular dysfunction in in both males and females *5x-FAD*. Remarkably, the mechanisms underlying this dysfunction show sex-specificity: In males, we observed lower BK_α_ expression and reduced Ca^2+^ sparks frequency; however, in females, there is an exacerbated and partially reversible nitro-oxidative modification of BK_α_.

## Methods

All animal procedures in this study were approved by the Institutional Animal Care and Use Committee of the University of Arizona College of Medicine (IACUC protocol 18-473) and are in accordance with the National Institutes of Health’s Guide for the Care and Use of Laboratory Animals, 8th edition. All animal experiments are reported in compliance with ARRIVE guidelines. The *5x-FAD* mice (5-6 months-old) used in this study are a transgenic model of early-onset AD that combines five different mutations in two genes associated with Familial AD. Three of the mutations are located in the gene that codes human amyloid precursor protein: Swedish (K670N, M671L), Florida (I716V), and London (V717I). The other two mutations are inserted in the gene of human presenilin, M146L and L286V. Transgene expression is controlled by the neuronal-specific *Thy1* promoter^29^. All mice were bred on a C57bl/6J background, and wild-type (WT) littermates were used as controls. Following euthanasia, the brain was collected and the pial arteries were dissected for patch clamp electrophysiology, pressure myography and quantification of vascular smooth muscle cells Ca^2+^ sparks activity by spinning disk confocal microscopy. The brain was either fixed in 4% paraformaldehyde for immunofluorescence, or flash-frozen for molecular biology and biochemical experiments (oxidized glutathione assay, affinity-purification of nitrosylated proteins followed by Western blot and qPCR). Baseline cortical perfusion and functional hyperemia following somatosensory stimulation were measured by laser speckle contrasting imaging.

Post-mortem brain tissues from CAA / AD and age-matched controls (no CAA or AD diagnosed) were used for assessment of BK_α_ S-NO by affinity-purification followed by Western blot. All human samples were kindly donated by the Brain and Body Donation Program from Banner Health, and are in compliance with all guidelines and requirements for patient de-identification. The complete and detailed methodology can be found in the *Supplemental Information - Methods*.

## Results

### Amyloid-β accumulation is higher in female 5x-FAD mice

One of the hallmarks of Alzheimer’s disease is the accumulation of amyloid-β peptides in the brain and cerebral microvasculature^30^. In this study, *5x-FAD* female mice showed a trend towards higher density of amyloid-β plaques in the cortex and hippocampus when compared with age-matched male *5x-FAD* (Figures S1A and S1B). No significant perivascular amyloid-β accumulation was observed in the parenchymal microcirculation of male or female *5x-FAD* mice (Figures S1C and S1D). We previously showed that pial arteries in this model exhibit extensive amyloid-β accumulation^31^. However, in our previous publication, we did not separate the incidence of pial vessel CAA by sex. Thus, we reanalyzed those data and observed no sex differences in pial vessel CAA in *5x-FAD* mice (Figures S1E, and S1F). Given the lack of microvascular intraparenchymal CAA in *5x-FAD*, we focused our vascular reactivity studies on the pial microcirculation.

### Increased contractility of pial arteries in 5x-FAD mice

Along with amyloid-β plaque accumulation, microvascular dysfunction is implicated in AD pathophysiology^30^. We observed an exacerbated contractile response in pial arteries of both female and male *5x-FAD* when compared to wild-type (WT) littermates, shown as higher spontaneous myogenic tone at a purported physiological intraluminal pressure compared to the respectively WT (50 mmHg, Figures 1A-B and 1E-F). Similarly, contractile response to a receptor-dependent stimulus (endothelin-1 [ET-1], 30 nM, Figures 1C and 1G) was higher in females and males *5x-FAD* mice than WT. No differences were observed in contractile responses to a receptor-independent depolarizing stimulus (60 mM [K^+^]_E_, Figures 1D and 1H). Thus, we conclude that contractile responses to a steady intraluminal pressure, as well as receptor-induced, are exacerbated in pial arteries of *5x-FAD* mice.

**Figure 1.**
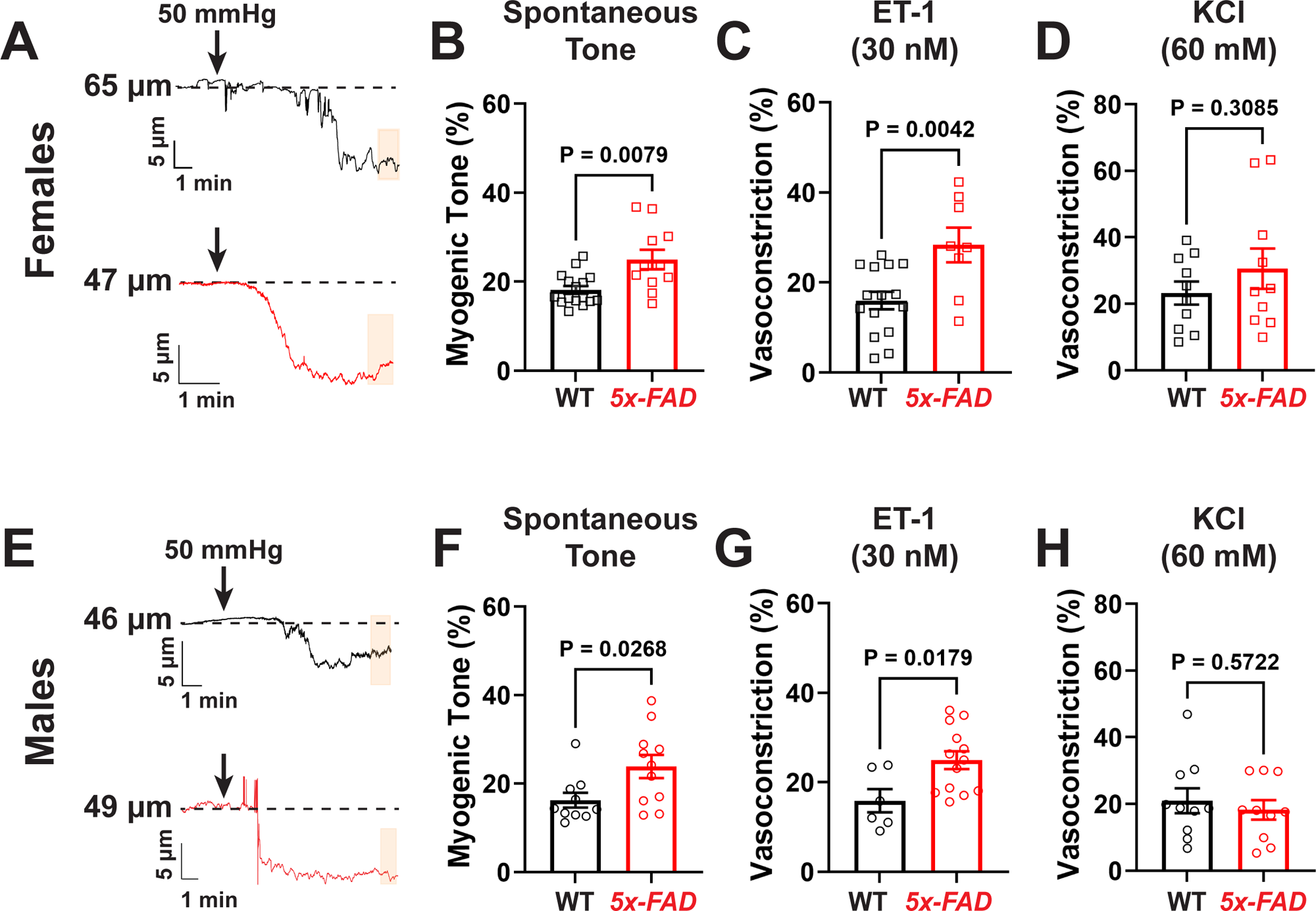
Cerebral microvascular hyper-contractility in 5x-FAD. **A,E)** Representative traces of the lumen diameter of pial arteries isolated from female (**A**) and male (**E**) WT (black, top) or *5x-FAD* (lower, red) during the generation of spontaneous myogenic tone. Arrows indicate the moment when intraluminal pressure was raised from 15 to 50 mmHg. **B, F)** Summary data showing higher spontaneous myogenic tone in pial arteries from female (**B**) and male (**F**) *5x-FAD*. **C, G)** Pial artery contractility to endothelin-1 (ET-1) was significantly increased in female (**C**) and male (**G**) *5x-FAD* when compared to WT littermates. **D, H)** Receptor-independent contraction to a depolarizing stimulus (60 mM KCl) was unchanged in female (**D**) and male (**H**). All data are means ± SEM. Each data point in graphs represents 1 pial artery from 1 mouse. Statistical analysis: unpaired two-tailed Mann-Whitney (**B, F**) or unpaired two-tailed *Student*’s t-test (**C, D, G, H**).

We then characterized autoregulatory responses (myogenic reactivity) of pial arteries in *5x-FAD* and WT littermates by step-wise increases in intraluminal pressure (from 5 mmHg to 160 mmHg). Under our experimental conditions, myogenic reactivity was not altered in *5x-FAD*, both females and males (Figure S2). Similarly, pial arteries from female or male *5x-FAD* do not undergo significant structural remodeling (Figure S3) or alterations in biomechanical properties (Figure S4). Thus, the enhanced contractility of pial arteries observed is independent of microvascular remodeling.

### Reduced BK_Ca_ channel function in pial arteries from 5x-FAD

BK_Ca_ channel is an important mechanism that provides negative feedback to myogenic constriction^32^. Therefore, to test whether the increased contractility observed in *5x-FAD* was due to BK_Ca_ dysfunction, we incubated pial arteries with the BK_Ca_ channel blocker iberiotoxin (IbTox, 30 nM). IbTox-induced vasoconstriction was significantly smaller in pial arteries isolated from *5x-FAD* when compared to WT littermates, for both females and males (Figures 2A-B and 2E-F), suggesting impaired BK_Ca_ activity in *5x-FAD* mice.

**Figure 2.**
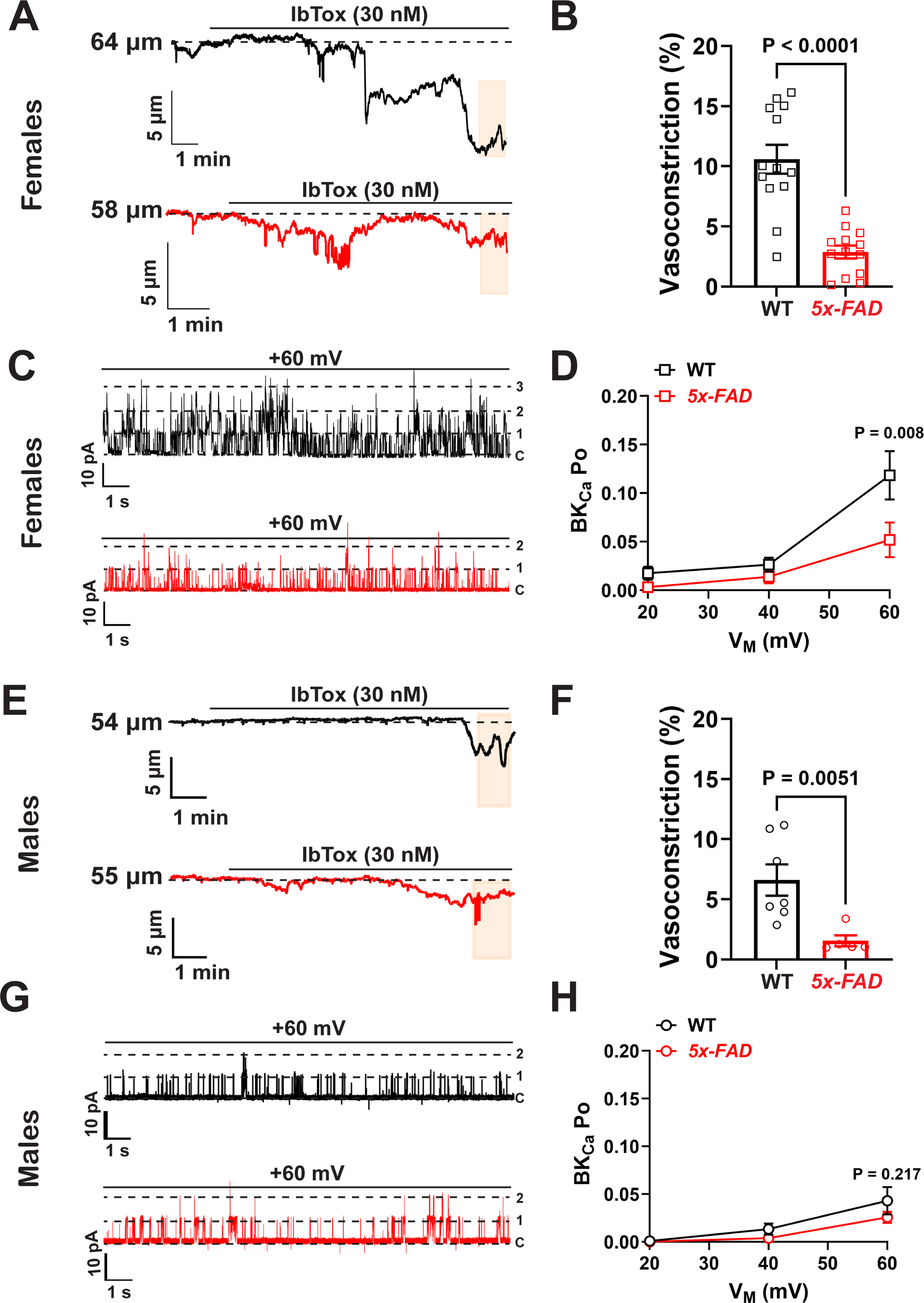
BK_Ca_ impairment in pial arteries from 5x-FAD. **A, E)** Representative traces of the lumen diameter of pial arteries isolated from female (**A**) and male (**E**) WT (black, top) or *5x-FAD* (lower, red) showing constriction to the BK_Ca_ blocker iberiotoxin (IbTox, 30 nM). Note the blunted constriction in *5x-FAD* mice. Orange boxes: regions where data were obtained**. B, F)** Summary bar graph showing blunted vasoconstriction to IbTox in female (**B**) and male (**F**) *5x-FAD*. Each data point in the graph represents 1 pial artery from 1 mouse. Statistical test: unpaired two-tailed Mann-Whitney test. **C, G)** Representative traces of an inside-out patch clamp from freshly-isolated pial artery smooth muscle cells showing single channel BK_Ca_ currents at a holding potential of +60 mV. Note the higher frequency of openings (downward deflections) in the patch from female (**C**) WT (top trace, black) compared to *5x-FAD* (lower trace, red). Single channel BK_Ca_ openings were not different between males WT and *5x-FAD* (**G**). C: closed; 1: 1 channel; 2: 2 channels. **D, H)** Summary graph showing BK_Ca_ open probability (Po) over a range of holding potentials (+20, +40, +60 mV). Po was significantly lower in inside-out patches from pial artery smooth muscle cells isolated from female *5x-FAD* mice (**D**), whereas BK_Ca_ Po was not different between males *5x-FAD* and WT littermates (**H**). Number of female replicates: WT n = 10 patches from 5 different mice; *5x-FAD* n = 7 patches from 4 different mice. Number of male replicates: WT n = 9 patches from 5 different mice; *5x-FAD* n = 9 patches from 6 different mice. All data are means ± SEM, two-way ANOVA with a Sidak *post-hoc* test.

We next corroborated the pressure myography findings by recording pial artery smooth muscle cell single channel BK_Ca_ currents in excised, inside-out patch-clamp electrophysiology experiments. We observed an approximately 50% lower BK_Ca_ open probability (Po) at +60 mV in patches from *5x-FAD* female than those from WT littermates (Figures 2C and 2D). However, in males, BK_Ca_ Po was not significantly different between WT and *5x-FAD* (Figures 2G and 2H).

Interestingly, when comparing IbTox constriction between WT females and males, we observed a significantly smaller constriction in pial arteries of males than in females (Vasoconstriction (%): 10.6 ± 1.2 vs 6.6 ± 1.3, WT females vs WT males, p = 0.0494, two-tailed *Student*’s t-test). This finding was again corroborated by patch-clamp electrophysiology: BK_Ca_ Po was significantly lower in WT males when compared to females (BK_Ca_ Po at +60 mV: 0.118 ± 0.025 vs 0.040 ± 0.013, WT females vs WT males, p = 0.0125, two-tailed *Student*’s t-test). These data suggest that physiological BK_Ca_ activity is lower in pial artery smooth muscle cells from males than in female WT mice.

### 5x-FAD males, but not females, have lower BK_α_ expression in pial arteries

To investigate if the reduced constriction to IbTox in *5x-FAD* mice was due to lower expression of BK_Ca_, we assessed mRNA expression of BK_α_ and BK_β1_ subunits in lysates of pial arteries from female and male *5x-FAD* and WT littermates by qPCR. Pial artery mRNA expression of both subunits was not significantly different between *5x-FAD* and WT in females (Figures 3A and S5A) or males (Figures 3C and S5B). Using Western blot, we then measured expression of the BK_α_ protein in pial artery lysates, and observed that, in females, expression of BK_α_ was similar between WT and *5x-FAD* (Figure 3B). However, expression of BK_α_ protein was significantly lower in male *5x-FAD* when compared to WT littermates (Figure 3D). An image of the membrane used for the inserts can be found in Figure S6.

**Figure 3.**
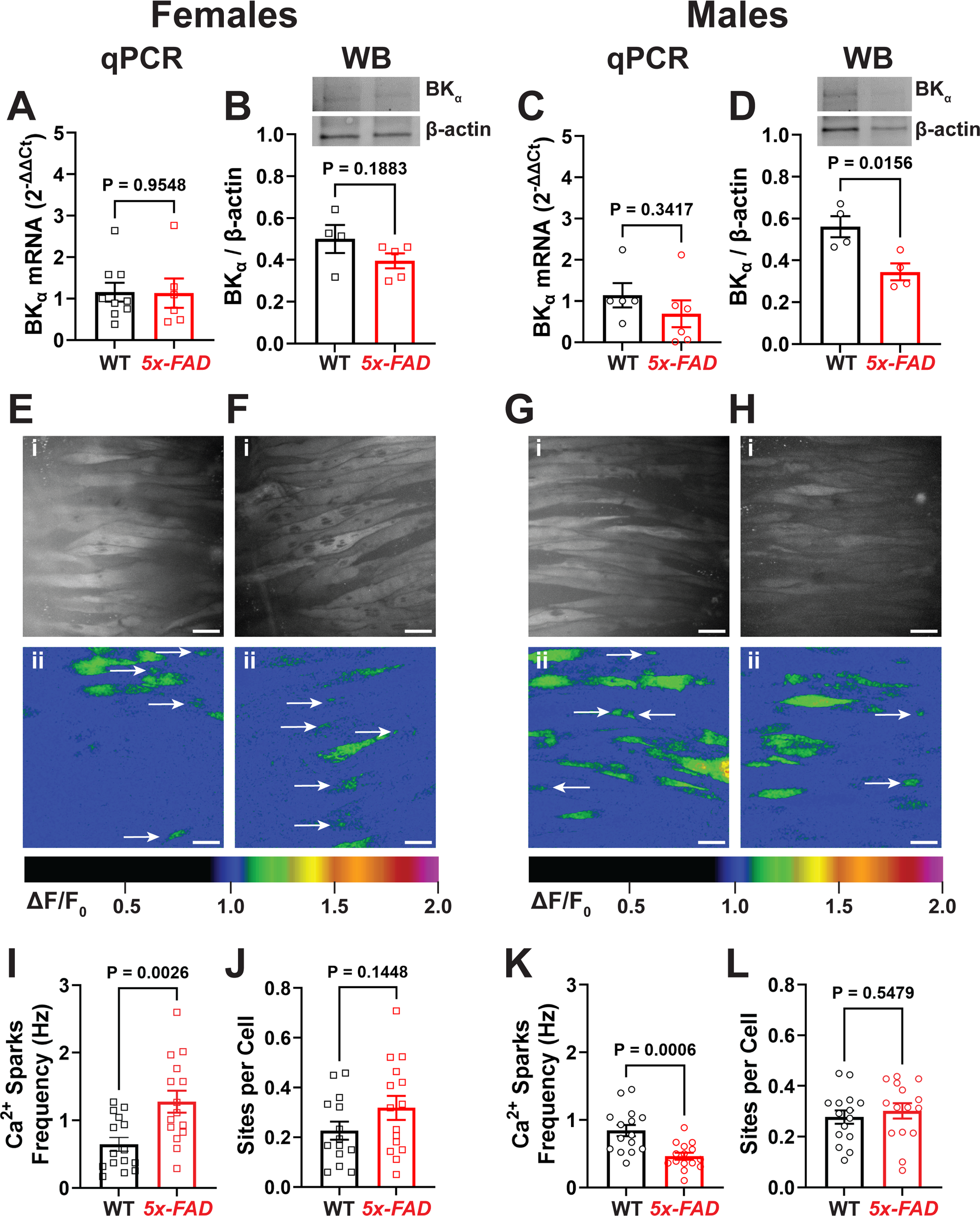
BK_Ca_ impairment in males is a consequence of lower BK_α_ expression and reduced Ca^2+^ sparks activity. **A-B)** Expression levels of BK_α_ mRNA (**A**) and protein (**B**) are not different between female *5x-FAD* and WT littermates. WB: Western blot. Each data point in the graph is a pial artery lysate from 1 mouse. Statistical analysis: unpaired two-tailed *Student*’s t-test. **C-D**) Expression levels of BK_α_ mRNA are similar between male *5x-FAD* and WT littermates (**C**); however BK_α_ protein expression is significantly lower in male *5x-FAD* when compared to WT littermates (**D**). Each data point in the graph is a pial artery lysate from 1 mouse. Statistical analysis: unpaired two-tailed *Student*’s t-test. **E-J)** Representative greyscale (**Ei, Fi**) and pseudocolored (**Eii, Fii**) images of fields of view of pial arteries from female WT (**E**) and *5x-FAD* (**F**) prepared *reverse en* face for recording of Ca^2+^ sparks (arrows). Ca^2+^ sparks frequency (**I**) was significantly higher in pial arteries from female *5x-FAD* than in WT littermates, without differences in the number of active sites per cell (**J**). Each data point in the graph is 1 field of view from a pial artery, for a total of 5 fields of view per artery from 3 different mice of each genotype. Statistical analysis: unpaired two-tailed *Student*’s t-test. **G-L)** Representative greyscale (**Gi, Hi**) and pseudocolored (**Gii, Hii**) images of fields of view of pial arteries from male WT (**G**) and *5x-FAD* (**H**) prepared *reverse en* face for recording of Ca^2+^ sparks (arrows). Ca^2+^ sparks frequency (**K**) was significantly lower in pial arteries from male *5x-FAD* than in WT littermates, without differences in the number of active sites per cell (**L**). Each data point in the graph is 1 field of view from a pial artery, for a total of 5 fields of view per artery from 3 different mice of each genotype. Statistical analysis: unpaired two-tailed *Student*’s t-test. All data are means ± SEM.

In brain lysates, we observed a downregulation of BK_α_ mRNA in female *5x-FAD* when compared to WT littermates (Figure S5C), without changes in mRNA for BK_β1_ (Figure S5D). No significant differences were observed in mRNA for BK_α_ and BK_β1_ in brain lysates from male *5x-FAD* when compared to WT (Figures S5E and S5F).

### Lower frequency of Ca^2+^ sparks in pial arteries from male 5x-FAD mice, but not in females

We next investigate whether the observed BK_Ca_ dysfunction was downstream from alterations in intracellular Ca^2+^ sparks, as shown previously in male APP23 mice^28^. Our data show that Ca^2+^ sparks frequency is significantly higher in pial artery smooth muscle cells of female *5x-FAD* mice than WT littermates (Figure 3I, Movies S1 and S2), without changes in the number of Ca^2+^ sparks sites per cell (Figures 3E, 3F and 3J). In contrast, Ca^2+^ spark frequency was significantly lower in pial artery smooth muscle cells of male *5x-FAD* than in WT (Figure 3K, Movies S3 and S4). This reduction was not due to a lower number of active Ca^2+^ sparks site per cell, as these were not different between males *5x-FAD* and WT (Figures 3G, 3H and 3L). Altogether, our findings show that BK_Ca_ dysfunction in *5x-FAD* male is due to lower expression of BK_α_ and reduced Ca^2+^ sparks frequency, whereas in females we observed an increase in Ca^2+^ sparks frequency without changes in BK_α_ expression.

### Oxidative stress impairs BK_Ca_ in female 5x-FAD

AD is associated with oxidative stress and inflammation that can contribute to vascular and neuronal damage^26,33^. Thus, we investigated if oxidative stress was present in the brain of *5x-FAD* using an oxidized glutathione assay. We observed that, in females, total glutathione (Total GSH) levels were similar between WT and *5x-FAD* (Figure 4A). However, brain levels of oxidized glutathione (GSSG, Figure 4B) and the ratio GSSG / Total GSH (Figure 4C) were significantly higher in female *5x-FAD* than WT littermates, suggesting the presence of an oxidative environment. None of the assessed parameters were altered in male *5x-FAD* when compared to WT littermates (Figure S7A-S7C).

**Figure 4.**
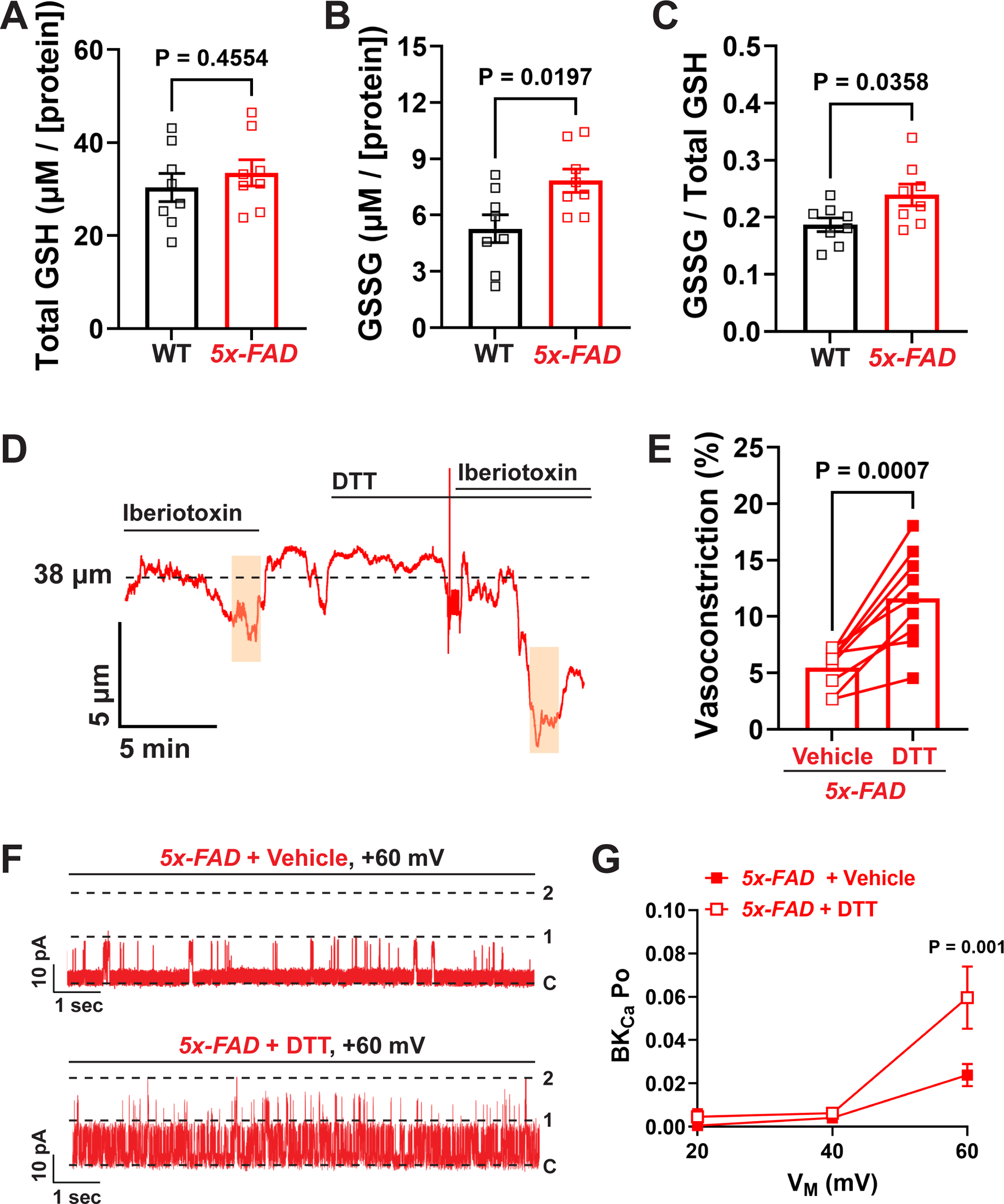
Oxidative-dependent BK_Ca_ impairment in pial arteries from female 5x-FAD. **A-C)** Glutathione assay from brain lysates of female *5x-FAD* showing the presence of an oxidative environment, observed as a significant increase in oxidized glutathione (GSSG, **B**) and the ratio between GSSG and reduced glutathione (GSH, **C**). Each data point represents a brain lysate from the cortex of an individual mouse. Statistical analysis: unpaired two-tailed Student’s t-test. D-E) Incubation of pial arteries from isolated female *5x-FAD* with the broad-spectrum reducing agent dithiothreitol (DTT, 10 µM) significantly recovers sensitivity to iberiotoxin (30 nM), as observed by the representative trace (**D**) and summary data (**E**). Orange boxes indicate the regions where data were extracted. Each data point in the graph represents 1 pial artery from an individual mouse. Statistical analysis: paired two-tailed *Student*’s t-test. **F-G)** Similarly, acute incubation with DTT partially recovers BK_Ca_ Po in freshly-isolated pial artery smooth muscle cell from female *5x-FAD*, as observed in the representative traces (**F**) and summary data (**G**). N = 16 patches from 6 different mice. Statistical analysis: Matching Mixed-Model Anova with a Sidak *post-hoc* correction for multiple comparisons. All data are means ± SEM.

We then hypothesized that this oxidant environment leads to reversible PTM in pial artery BK_Ca_ in female *5x-FAD*, thus causing its impairment. We tested this hypothesis by incubating *ex vivo* pial arteries and excised inside-out membrane patches with the broad-spectrum reducing agent dithiothreitol (DTT, 10 µM). We observed that DTT partially reversed pial artery sensitivity to IbTox in females, observed as a larger vasoconstriction response after incubation with DTT (Figures 4D and 4E). These findings were corroborated in patch-clamp electrophysiology experiments, as incubation of patches with DTT partially restored BK_Ca_ Po (Figures 4F and 4G). DTT did not affect BK_Ca_ Po in pial artery smooth muscle cells isolated from male *5x-FAD* (Figure S7D-S7E). Altogether, these data suggest that the oxidative environment in female, but not males, *5x-FAD* leads to oxidation of BK_Ca_, consequently reducing its function in pial artery smooth muscle cells.

### Exacerbated BK_Ca_ S-nitrosylation in female 5x-FAD and CAA / AD patients

Next, we sought to identify the likely type of PTM responsible for the observed BK_Ca_ impairment. The inducible nitric oxide synthase (iNOS) isoform is a pro-inflammatory enzyme that is increased in the AD brain^34,35^. This isoform produces a high amount of NO, which can lead to nitro-oxidative damage of cellular structures^36,37^, including excessive nitrosylation of target proteins. Thus, we investigated if iNOS expression is increased in the pial arteries of *5x-FAD* females. Using qPCR, we observed that the expression of iNOS mRNA was increased in the pial arteries from female *5x-FAD* mice compared to WT (Figure 5A). No significant differences were observed for endothelial (eNOS, Figure S8A) or neuronal NOS (nNOS, Figure S8B) in pial arteries. To corroborate the mRNA findings, we performed a semi-quantitative assessment of fluorescence intensity of iNOS immunolabeling in pial arteries, which was significantly higher in pial vessels from female *5x-FAD* than in WT mice (Figures 5B and 5C). These results suggest that iNOS is upregulated on the pial vessels from female, but not male, *5x-FAD* mice. Lastly, inhibition of NOS with L-NAME (200 µM) induced a greater pial artery constriction in *5x-FAD* than in WT littermates (Figure S8C).

**Figure 5.**
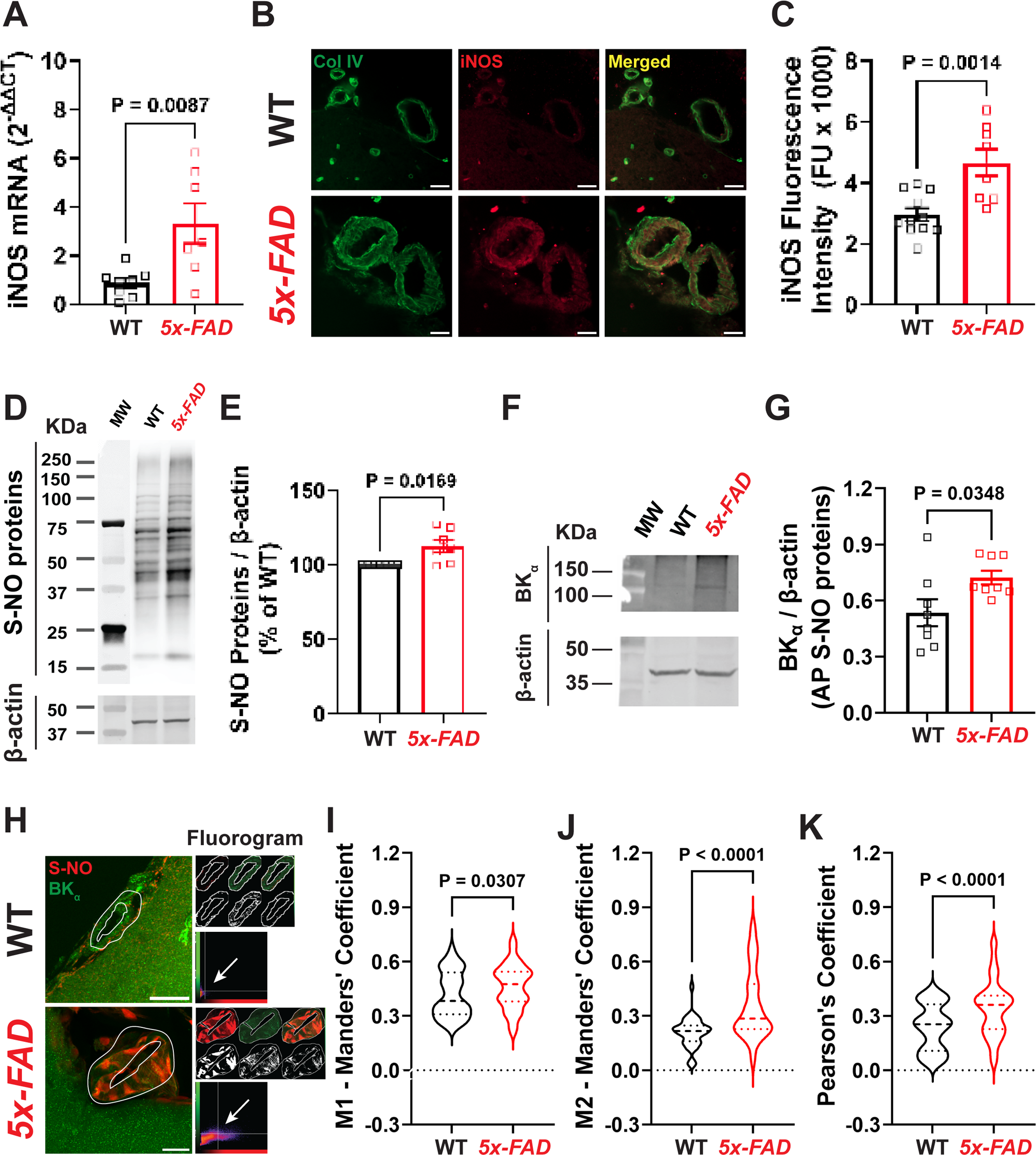
Exacerbated BK_Ca_ S-NO in female 5x-FAD. **A)** Higher iNOS mRNA expression in lysates of pial arteries isolated from female *5x-FAD* than in WT littermates. Each data point represents 1 pial artery lysate from an individual mouse. Statistical analysis: unpaired two-tailed *Student*’s t-test. B) Immunofluorescence labeling of iNOS (red) in pial arteries (collagen IV – Col IV, green). Note the apparent higher fluorescence intensity in iNOS labeling in pial arteries from female *5x-FAD* when compared to WT littermates. Bar = 20 µm. **C)** Blinded, randomized semi-quantitative analysis of iNOS fluorescence intensity in pial arteries of female WT and *5x-FAD*. Each data point represents the average fluorescence intensity of 5 pial arteries randomly imaged from an individual mouse. Statistical analysis: unpaired two-tailed *Student*’s t-test. **D-E)** Quantification of global S-NO proteins in cortical lysates of *5x-FAD* and WT littermates. Note the higher expression of S-NO protein in lysates from *5x-FAD* cortex when compared to WT in the representative blot (**D**) and summary data (**E**). Each data point represents a lysate from an individual mouse. Statistical analysis: unpaired two-tailed Mann-Whitney test. **F-G)** Affinity-purification (AP) of S-NO proteins in brain lysates followed by Western blot against BK_α_ show a significantly higher expression of BK_α_ S-NO in female *5x-FAD* when compared to WT littermates, as evidenced by the representative blot (**F**) and summary data (**G**). Each data point represents a lysate from an individual mouse. Statistical analysis: unpaired two-tailed *Studen*t’s test. **H-K)** Co-localization analysis using Mander’s M1 (**I**) and M2 (**J**) coefficients, as well as the Pearson’s coefficient (**K**) show a significantly higher co-localization between of BK_α_ (**H**, green) and global S-NO proteins (**H**, red) in pial arteries from female *5x-FAD* (fluorogram in **H**) Bar = 20 µm. Violin plots represent the distribution of co-localized pixels in each image. Imaging of pial arteries was performed in a blinded and randomized manner, a total of 3 arteries were imaged per mouse for a total of 5 mice per group. Statistical analysis: unpaired two-tailed Student’s t-test. Data in violin plots are median ± 95% confidence intervals, all other data are means ± SEM.

Excessive NO production in the presence of an oxidative environment can lead to excessive protein S-NO^38^, thus we investigated if this particular PTM was involved in the BK_Ca_ dysfunction observed in female *5x-FAD*. Our findings show that total S-NO proteins were significantly higher in brain lysates of female *5x-FAD* when compared to WT littermates (Figures 5D-5E), but not in male *5x-FAD*, where they were actually significantly reduced (Figure S9). We then performed an affinity-purification of S-NO proteins from brain lysates of female *5x-FAD* and WT littermates, followed by Western blot against the BK_α_ subunit. We observed a significant increase in the amount of BK_α_ S-NO (Figures 5F-G, whole blot of inserts in Figure S10). We corroborated these findings by performing a co-immuno-localization assay to investigate vascular-specific BK_α_ S-NO (Figure 5H). Our data shows that there is a significant increase in co-localization between the BK_α_ subunit and S-NO proteins, as evidenced by the M1 / M2 – Mander’s coefficients (Figures 5I-5J) and the Pearson’s coefficient (Figure 5K). Together, these findings support our hypothesis that the pro-nitro-oxidative environment in *5x-FAD* females leads to excessive BK_α_ S-NO.

We then investigated if the increase in BK_α_ S-NO observed in *5x-FAD* was also present in humans with CAA or AD. We obtained post-mortem brain samples from the Banner Brain and Body Donation Program, from age-matched women and men, and separated them in 3 groups: healthy (non-CAA / non-ADA, n=6), CAA / non-AD (CAA, n=6) and AD + CAA (n=12). Healthy patients were defined no presence of dementia or symptoms of parkinsonism, and no presence of CAA or senile plaques in histopathology. CAA / non-AD were patients without dementia or parkinsonism, but with evident perivascular amyloid plaques and no presence of senile plaques. AD + CAA were patients with Alzheimer’s disease with a minimum of intermediate or high NIA-Reagan criteria^39^, and presence of CAA and / or senile plaques. Using Western blot, we observed that there was a trend towards an increase in BK_α_ subunit expression in brain from AD + CAA patients (p=0.0976, Brown-Forsythe and Welch’s ANOVA), without evident sex differences (Figures 6A-6B). When performing an affinity purification of S-NO proteins followed by Western blot, we again observed a trend towards an increase in BK_α_ S-NO in AD + CAA patients (p=0.0772, Brown-Forsythe and Welch’s ANOVA), without sex differences (Figures 6C-6D). Images of the whole blots shown in the cropped insets (Figures 6A and 6C) can be found in the Figure S11. These findings suggest that BK_α_ S-NO is also present in AD patients, although in both women and men.

**Figure 6.**
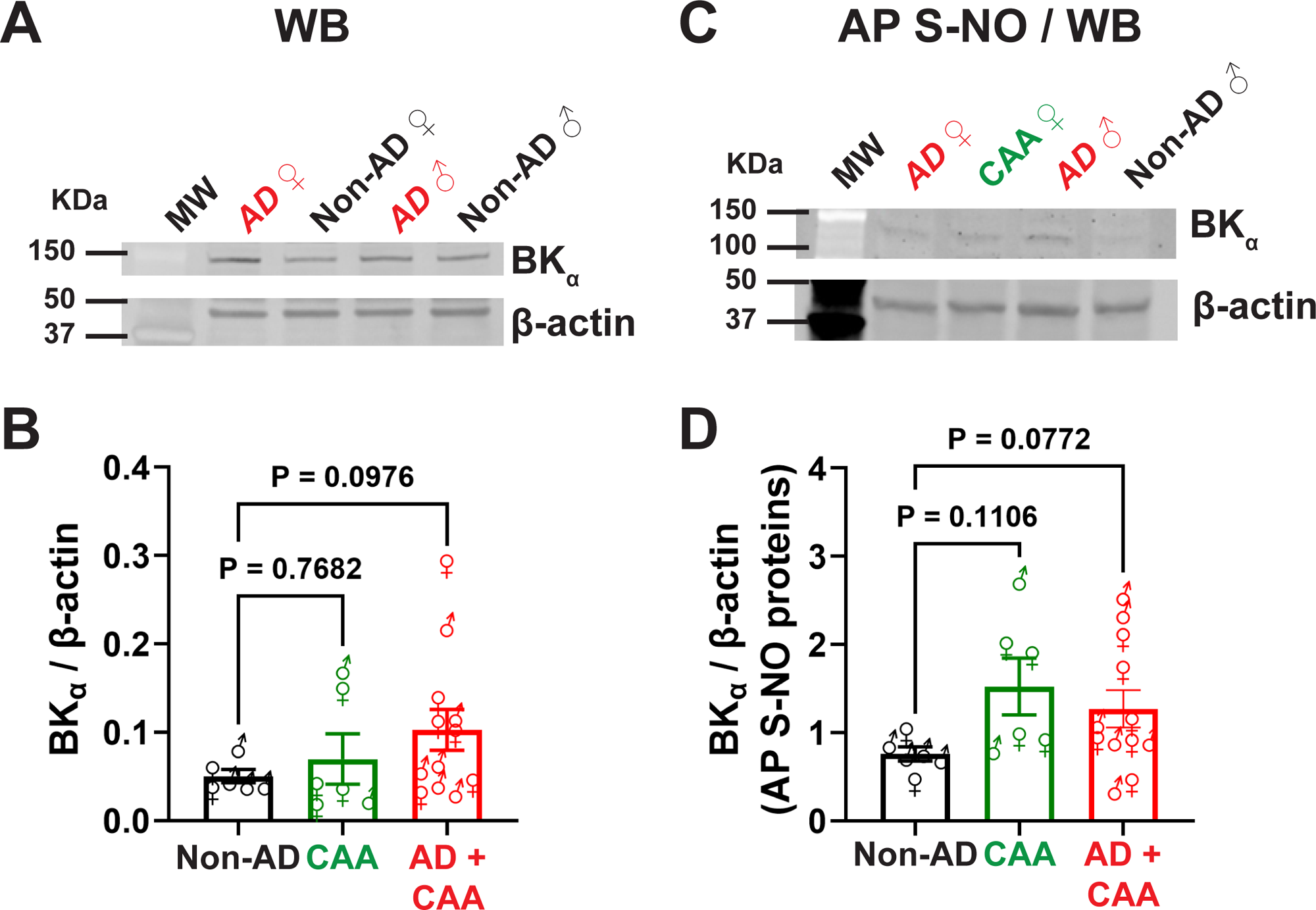
BK_Ca_ S-NO in post-mortem human brain samples. **A-B)** Western blot (WB) quantification of BK_α_ expression in *post-mortem* brain lysates of patients without CAA / AD (Non-AD), CAA or AD + CAA. Representative blots are shown in **A** and summary data in **B**. Each data point represents a brain lysate from an individual age-matched woman (♀) or man (♂). Statistical analysis: Brown-Forsythe and Welch ANOVA with a Dunnett T3 correction for multiple comparisons. **C-D)** Affinity-purification (AP) of S-NO proteins in brain lysates followed by WB against the BK_α_ subunit show a trend towards an increase in BK_α_ S-NO in AD + CAA patients when compared to non-AD, as evidenced by the representative blots in **C** and summary data in **D**. Each data point represents a brain lysate from an individual age-matched woman (♀) or man (♂). Statistical analysis: Brown-Forsythe and Welch ANOVA with a Dunnett T3 correction for multiple comparisons. All data are means ± SEM.

### Cerebral perfusion deficits in female 5x-FAD

Next, we investigated if there are changes in basal cortical perfusion in *5x-FAD* females and males compared to sex-matched WT littermates. Using laser speckle contrast imaging, we observed that the baseline perfusion of the entire superficial cortex was not different between WT and *5x-FAD* females (Figures S12B-S12C, WC: whole cortex) or males (Figures S12D-S12E, WC: whole cortex). However, when analyzing discrete perfusion of specific vascular territories (diagram in Figures S12A)^40^, we observed that *5x-FAD* females showed lower basal perfusion to the frontal cortex (Figures S12B-S12C), but not in other regions. No significant differences were observed in cortical perfusion of male *5x-FAD* mice (Figures S12D-S12E).

### Impaired functional neurovascular responses in females and males 5x-FAD

We next tested the hypothesis that impaired vascular BK_Ca_ would result in neurovascular dysfunction in *5x-FAD*. First, we showed that acutely inhibiting BK_Ca_, achieved by applying IbTox (30 nM) atop the thinned-skull cranial window, significantly blunted hyperemic responses on the somatosensory cortex during mechanical stimulation of contralateral whiskers^41^ (Figures 7A-7C), suggesting that BK_Ca_ play a role in neurovascular responses. We then assessed neurovascular responses in *5x-FAD* mice and compared to WT littermates. We observed that both females (Figures 7D-7F) and males (Figures 7G-7I) *5x-FAD* had significantly lower hyperemic responses than WT littermates. Together, these findings suggest that the cerebral vascular impairment observed in *5x-FAD* mediates neurovascular dysfunction in the model.

**Figure 7.**
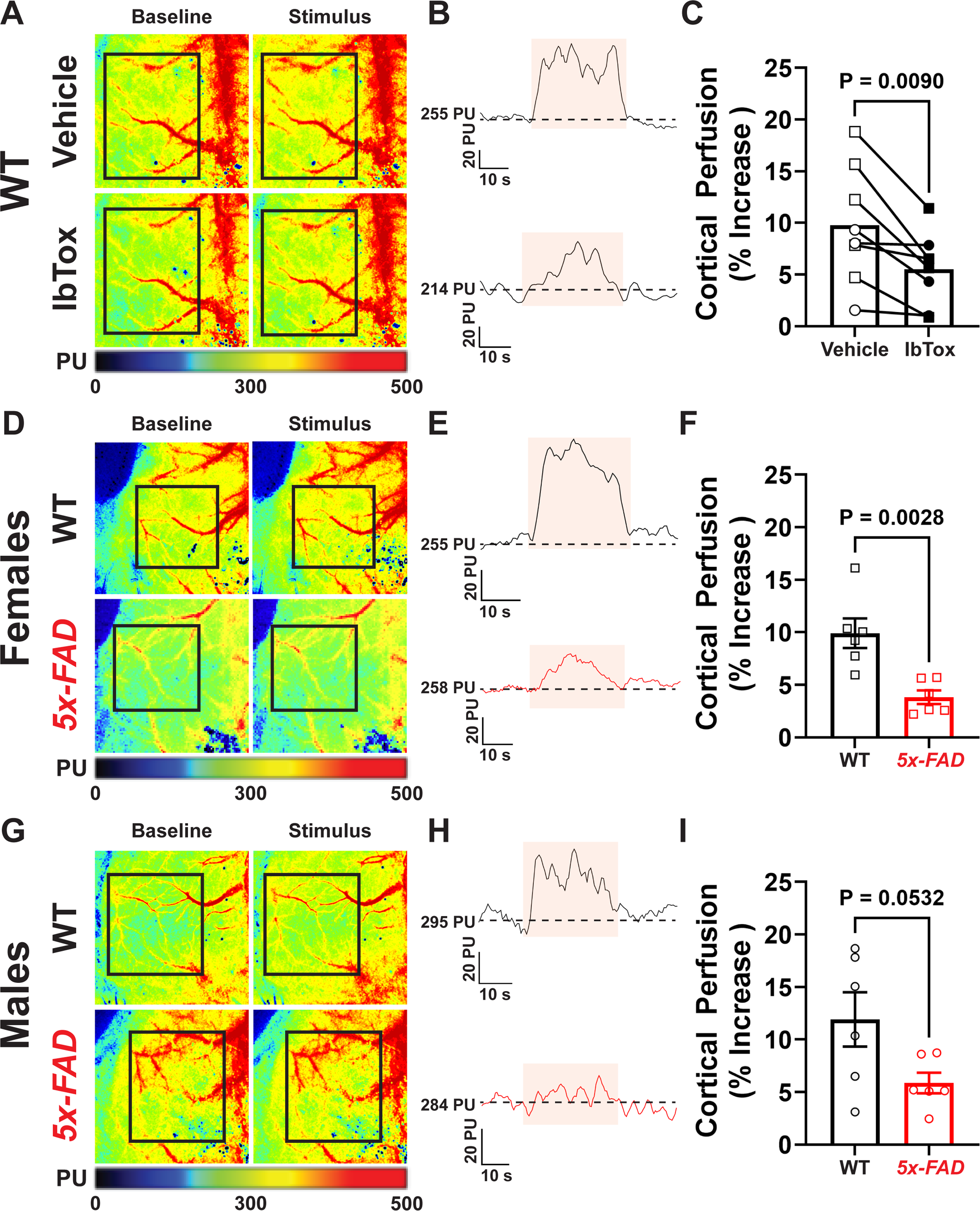
Impaired neurovascular coupling after acute BK_Ca_ inhibition and in 5x-FAD. **A-C)** Acute application of the BK_Ca_ blocker iberiotoxin (IbTox, 30 nM) atop the thinned-skull cranial window blunted neurovascular responses following somatosensory stimulation in WT mice, as evidenced by the pseudocolored perfusion maps (**A**), representative traces (**B**) of perfusion within the region of interest (black rectangle) and summary data (**C**). Each data point represents an individual mouse, females in square (□) and males in circles (○), after application of vehicle (□, ○) or IbTox (■,●). Statistical analysis: paired two-tailed *Student*’s t-test. **D-I)** Neurovascular coupling responses were significantly impaired in female (**D-F**), and showed a trend towards decrease in male (**G-I**), *5x-FAD* when compared to WT littermates, as evidenced by the perfusion maps (**D, G**), representative perfusion traces (**E, H**) and summary data (**F, I**). Each data point represents an individual mouse. Statistical analysis: paired two-tailed *Student*’s t-test. PU = perfusion units. All data are means ± SEM.

## Discussion

Our findings show that cerebral BK_Ca_ microvascular dysfunction is present in both female and male *5x-FAD*, likely mediating neurovascular impairments in the model. A major novelty of our study lies in our findings elucidating the distinct mechanisms underlying BK_Ca_ dysfunction in females and males. In males, we observed reduced expression of the BK_α_ subunit and lower Ca^2+^ sparks frequency. On the other hand, in females, we show that BK_α_ expression is unchanged and Ca^2+^ sparks frequency is higher, yet BK_Ca_ activity is likewise impaired. Upon further investigation, we uncovered that the BK_α_ subunit in female *5x-FAD* undergoes oxidative modification, particularly excessive S-NO, and this partially underlies cerebral microvascular BK_Ca_ dysfunction. The exacerbation in nitrosylation is downstream from increased vascular iNOS expression and general oxidative stress in the brain environment, none of which were present in males (Figure S13). Lastly, we describe two other novel findings in our study: basal cerebral microvascular BK_Ca_ activity seems lower in male than in female C57Bl/6J mice, and that BK_Ca_ function is involved in physiological neurovascular responses, as its blockade blunts somatosensory-induced functional hyperemia. Altogether, our data highlight the complexities of microvascular regulation in the context of AD, and the differential effects of biological sex that may hinder putative therapies ineffective as a “one size fits all” approach.

AD is a progressive neurodegenerative disease characterized by accumulation of amyloid-β and neurofibrillary tangles, particularly in the temporal lobe and neocortical structures^42^, which lead to cognitive deficits^30^. Beyond misfolded protein pathology, cerebral vascular dysfunction contributes to AD progression, and much attention has been placed on loss of endothelial function and blood-brain-barrier breakdown^43,44^. However, less is known on how cerebral microvascular smooth muscle cell function is affected in AD, as well as its participation in AD’s development and progression. A recent report by Taylor *et al.* described that acute exposure of cerebral arteries isolated from male C57Bl/6 mice to amyloid-β_(1-40)_ increases myogenic tone due to impaired BK_Ca_ activity^28^. Our findings expand on this report, and we show that not only spontaneous myogenic tone is higher in *5x-FAD*, a mouse model of AD with accumulation of both amyloid-β_(1-40)_ and amyloid-β_(1-42)_ (predominant), but so is contractility to ET-1. These data suggest the presence of a general hyper-contractile state in the cerebral microcirculation in both female and male arteries during AD progression. Our data suggests that BK_Ca_ impairment likely underlies this hyper-contractile status in *5x-FAD*, as its function is impaired in *ex vivo* pressurized arteries. Paradoxically, we did not observe any significant changes in autoregulation. Although loss of autoregulation is observed in cerebral small vessel disease^45^ and it was shown to occur in some rodent models of AD^46^, human studies have also failed to observe cerebral autoregulation impairments in AD patients^47^. The mechanistic explanation underlying these differences in autoregulation between AD models and humans remain unclear and require further exploration.

Impairment in BK_Ca_ function is associated with vascular dysfunction in various disease models, including diabetes^48,49^, hypertension^50,51^, menopause^41,52^, cigarette smoking^53^ and AD^28^. One limitation of the aforementioned studies is that they were all performed in a single sex, and the vast majority in males (with the exception of studies focused on menopause). Our findings expand on these previous observations and show a sex-dependent mechanism of BK_Ca_ impairment in *5x-FAD*. We show that BK_Ca_ activity is reduced in pial arteries independently of biological sex, observed as blunted constriction to iberiotoxin. However, the cellular and molecular mechanisms responsible for the lower BK_C_a activity diverge between females and males. We observed that, in male *5x-FAD*, single channel BK_Ca_ Po was not lower than that recorded from WT littermates, indicating that BK_Ca_ dysfunction was likely upstream from intracellular signals regulating BK_Ca_ activity. Upon further investigation, we describe that expression of the pore-forming subunit BK_α_ is lower in male *5x-FAD* than in WT littermates, which could account for lower sensitivity to Iberiotoxin in pressure myography experiments. In addition, frequency of Ca^2+^ sparks was significantly reduced in male *5x-FAD*, similar to data published by Taylor *et al.* in male APP23 mice^28^. On the other hand, female *5x-FAD* showed reduced single channel BK_Ca_ Po despite similar levels of BK_α_ expression and an increase in Ca^2+^ spark frequency, suggesting that, in females, BK_Ca_ dysfunction was likely due to molecular alterations in the BK_α_ subunit itself, such as PTMs resulting from nitro-oxidative stress.

In support of this hypothesis, we observed that brains of females *5x-FAD*, but not males, show an increase in markers of oxidative and nitrosative stress, observed as higher levels of oxidized glutathione (GSSG) and higher levels of global S-NO proteins. Together, this nitro-oxidative environment can result in exacerbated PTMs within the BK_α_ subunit. PTMs are the result of binding of different moieties to proteins, such as phosphorylation, acetylation, lipidation, oxidation and nitrosylation^24,54^. PTMs play a vital role in the known heterogeneity and tissue specificity of the biophysical properties of BK_Ca_, as they have profound effects on gating, voltage and Ca^2+^ sensitivity, and membrane availability of the channel^16^. However, how S-NO affects BK_Ca_ remains controversial: some studies show that NO can, directly or indirectly, activate BK ^55,56^, whereas others show that NO itself, or peroxynitrite, do not affect or can blunt BK_Ca_ activity. For instance, a NO-donor failed to activate the BK_Ca_ channel in a heterologous expression system^57^. Moreover, Liu *et al.* showed that peroxynitrite reduces the open-state probability of BK_Ca_ channels in isolated vascular smooth muscle cells^58^. We show that, in female *5x-FAD*, the BK_α_ subunit undergoes excessive S-NO, and this nitro-oxidative PTM likely mediates channel dysfunction in a reversible manner. This assertion is supported by our data that the broad-spectrum reducing agent DTT partially recover single BK_Ca_ Po and arterial responsiveness to iberiotoxin. Although the molecular mechanisms of how S-NO impair BK_Ca_ remain undetermined, we speculate that they may reduce Ca^2+^ sensitivity of the BK_α_ subunit, which may explain the increase in Ca^2+^ sparks frequency observed in female *5x-FAD*, likely to compensate for such lower sensitivity. This, however, remains untested.

We further made efforts to elucidate the cellular source responsible for excessive protein S-NO, and observed an increase in cerebral vascular iNOS expression. Inducible NOS (NOS2) is commonly upregulated under stress and inflammatory conditions^34^, such as is observed in AD and other dementias^33,59^. Distinctly from the other NOS isoforms (eNOS and nNOS), iNOS is effectively locked in an active position due to the tight binding of calmodulin^34,60^, leading to sustained basal NO production. Together with oxidative stress, the higher NO production can result in excessive protein S-NO, as we observed in female *5x-FAD*. Further, we observed a trend towards an increase in BK_α_ S-NO in *post-mortem* brain samples from patients with AD + CAA when compared to age-matched non-dementia controls (healthy), without any evident changes in BK_α_ expression. Interestingly, in humans, our data does not indicate the presence of sex differences, as both women and men show BK_α_ S-NO. The discrepancies between the human and *5x-FAD* findings are not immediately apparent, but we speculate that other co-morbidities in males may increase BK_α_ S-NO in men, including cigarette smoking or cardiovascular disease. We currently do not have the medical records of the patients included in our samples; therefore, this possibility remains a speculation.

Ultimately, our observed microvascular BK_Ca_ dysfunction may underlie the functional perfusion deficits observed in *5x-FAD*. We show that acute application of iberiotoxin atop the cranial window blunts neurovascular responses in both female and male C57Bl/6J mice, showing the role of BK_Ca_ in these functional hemodynamic responses, as shown by others using an *ex vivo* brain slice preparations^61^. Further, both females and males *5x-FAD* mice showed a reduction in neurovascular responses during somatosensory stimulation. A similar neurovascular impairment was recently reported by us, in which female mice that underwent chemically-induced menopause showed reduced functional hyperemia responses associated with cerebral microvascular BK_Ca_ impairment^41^. However, it is important to note that these studies, and our data presented herein, are associative, and further experiments are needed to definitively show the role of cerebral microvascular BK_Ca_ in neurovascular coupling responses. Regardless, our previous and current findings, in conjunction with findings from others^28,51,62^, reiterate the central role played by BK_Ca_ in basal physiological hemodynamic regulation in the brain, as well as its importance in cerebrovascular diseases.

Lastly, the molecular basis of the sex differences observed remains unclear. Sex differences are commonly reported in mouse models of AD^63,64^, as well as in humans, and sex hormones seem to play a role. Female *5x-FAD* and AD patients have lower expression of aromatase in the hippocampus, an important enzyme that produces 17β-estradiol^65^. Since estradiol can play a role in vascular function^66,67^, neuroplasticity^68^, synaptic stability^68^, as well as attenuation of oxidative stress^69^, it is possible that reduction in its concentration or bioavailability may result underlie the heightened AD pathology in females. Further, in *5x-FAD* mice harboring human ApoE3 or ApoE4, ovariectomy worsens cognitive function, and β-estradiol partially recovers cognition in *5x-FAD* x ApoE3, with little effect in *5x-FAD* x ApoE4. Testes levels of testosterone are higher in male *5x-FAD* than in wild-type, without changes in sperm viability, testosterone levels are higher^70^. Currently, there are no published studies showing the effects of castration on *5x-FAD* males.

In conclusion, our results show that the BK_Ca_-dependent microvascular and neurovascular dysfunction occurs in both male and female *5x-FAD*, although the mechanisms underlying BK_Ca_ impairment differ based on biological sex. Further, we show that one of such mechanisms, namely excessive BK_α_ S-NO, may also be present in humans with AD and CAA, suggesting that manipulation of this basic pathophysiological mechanism could have therapeutic potential.

### Novelty and Significance

Cerebrovascular dysfunction is a hallmark of Alzheimer’s disease and related dementias. A property of the resistance vasculature is to constrict when pressurized (myogenic tone); however, over-constriction can be detrimental. Feedback mechanisms, including the opening of BK_Ca_, exist to keep the system in balance. Here, using a combination of molecular biology with *ex vivo* and *in vivo* assessments, we show a novel mechanism associated with BK_Ca_ dysfunction in the cerebral microvasculature of female *5x-FAD* mice. We report increased BK_Ca_ S-nitrosylation linked to higher myogenic tone and reduced channel activity. These changes were associated with lower perfusion of the frontal cortex and impaired neurovascular reactivity, suggesting that nitro-oxidative stress is an important mechanism of vascular dysfunction in Alzheimer’s disease in mice and human.

## Supporting information

SI movie 5xFAD female

SI movie 5xFAD male

SI movie WT female

SI movie WT male

## Acknowledgments

We thank Patricia Jansma, co-manager of the Office Research, Innovation & Impact’s Imaging Cores - Optical Facility at the University of Arizona, for providing training service at the microscope and helping with the microscopy image acquisition. We would further like to thank Dr. Anne M. Dorrance (Michigan State University) for her critical review of our manuscript.

We are grateful to the Banner Sun Health Research Institute Brain and Body Donation Program of Sun City, Arizona for the provision of human biological materials. The Brain and Body Donation Program has been supported by the National Institute of Neurological Disorders and Stroke (U24 NS072026 National Brain and Tissue Resource for Parkinson’s Disease and Related Disorders), the National Institute on Aging (P30 AG19610 and P30AG072980, Arizona Alzheimer’s Disease Center) the Arizona Department of Health Services (contract 211002, Arizona Alzheimer’s Research Center), the Arizona Biomedical Research Commission (contracts 4001, 0011, 05-901, and 1001 to the Arizona Parkinson-s Disease Consortium) and the Michael J. Fox Foundation for Parkinson’s Research.

The mouse strain used for this research project, B6SJL-Tg(APPSwFlLon, PSEN1*M146L *L286V)6799Vas/Mmjax, RRID:MMRRC_034840-JAX, was obtained from the Mutant Mouse Resource and Research Center (MMRRC) at The Jackson Laboratory, an NIH-funded strain repository, and was donated to the MMRRC by Robert Vassar, Ph.D., Northwestern University.

## Supplemental Material

A detailed description of the materials and methods is provided in the *Supplemental Material*.

## Author Contribution

**JFS** – conceived, designed, performed the experiments, analyzed data, and wrote the initial draft of the manuscript;

**FDP** – performed experiments and analyzed data;

**PM –** performed experiments and analyzed data;

**SHT –** performed experiments and analyzed data;

**ECL –** performed experiments and analyzed data;

**AS** – performed the experiments and analyzed data;

**AMK** – performed experiments and analyzed data;

**MTG** – performed the experiments and analyzed data;

**PWP** – conceived, designed, performed experiments, analyzed data and revised the draft of the manuscript.

All authors declare that they reviewed and edited the manuscript, and agree to its submission.

