## Supplementary material for "Sex-specific mechanisms of cerebral microvascular BK_Ca_ dysfunction in a mouse model of Alzheimer’s disease": SI Appendix Methods

### Supplementary information

### Methods

#### Animals

Five to six (5-6) month-old male and female *5x-FAD* mice were used in this study. *5x-FAD* transgenic mice were originally purchased from Jackson Laboratories (strain number 34840-JAX) and are currently maintained in our breeding colony at the University of Arizona. They are a transgenic model of early-onset AD that combines five different mutations in two genes associated with Familial AD. Three of the mutations are located in the gene that codes human amyloid precursor protein: Swedish (K670N, M671L), Florida (I716V), and London (V717I). The other two mutations are inserted in the gene of human presenilin, M146L and L286V. Transgene expression is controlled by the neuronal-specific *Thy1* promoter[1]. All mice were bred on a C57bl/6J background, and wild-type (WT) littermates were used as controls. Mice were housed at the animal facilities at the University of Arizona, 4 mice per cage (20.7 x 31.6 x 21.5 cm), in microisolated racks with access to water and food *ad libitum*, and light-dark cycles of 12 hours each and room temperature of 25 ± 1°C.

All animal procedures in this study were approved by the Institutional Animal Care and Use Committee of the University of Arizona College of Medicine (IACUC protocol 18-473) and are in accordance with the National Institutes of Health's Guide for the Care and Use of Laboratory Animals, 8th edition. All animal experiments are reported in compliance with ARRIVE guidelines.

#### Amyloid-β plaque staining

For amyloid-β plaque accumulation, brain sections (50 µm-thick) from perfusion-fixed mice were permeabilized in 0.1% Triton X-100 in PBS overnight, then treated with 15% formic acid in PBS for 15 minutes to enhance amyloid-β staining, and subsequently washed with PBS. Then, they were incubated in a 0.5% thioflavin-T solution for 10 minutes, protected from light, and subsequently washed with 80% ethanol, 95% ethanol, and then PBS prior to mounting onto a slide. Brain sections were then washed and mounted on glass slides using Fluoroshield mounting media. Images were acquired and stitched using SoftWoRx v1.2 (Applied Precision Inc., Issaquah, WA) on a DeltaVision Core system (GE Healthcare Biosciences, Piscataway, NJ) consisting of an Olympus IX71 microscope equipped with a CoolSNAP HQ2 camera (Teledyne Photometrics, Tucson, AZ) and a 10x objective. Separate images of the brain were acquired and stitched together to generate a single coronal section of the brain. Amyloid-β plaque burden in the cortex and hippocampus were assessed by manually counting the number of plaques in each region then dividing by the total area of the region using ImageJ. Data are shown as amyloid-β plaque density (number / $mm^2$ of tissue).

**Assessment of intraparenchymal and pial vessel CAA**

Brain sections (50 μm-thick) of perfusion-fixed female and male *5x-FAD* were used for immunolabeling to quantify CAA, as previously described[2]. All immunofluorescence labeling in this study followed the same initial protocol: mice were euthanized using isoflurane (5% mixed with breathing air) for cardiac perfusion-fixation with 4% paraformaldehyde. Brains were collected and post-fixed in 4% paraformaldehyde for 24 hours at room temperature, then washed with PBS. Brains were sectioned using a Leica VT1000P vibratome (Leica Biosystems Nussloch GmbH, Nußloch, Germany) to obtain 50 μm thick coronal slices. Brain slices were permeabilized in 0.1% Triton X-100 in PBS overnight and treated with 0.5% sodium borohydride for 15 minutes to reduce the tissue autofluorescence and washed with PBS. After washes, brain sections were blocked using 5% horse serum, 0.1% Triton X-100 in PBS for 1 hour, then incubated with primary antibodies. For intraparenchymal CAA, free-floating brain slices were co-immunolabeled with antibodies diluted in 5% horse serum, 0.1% Triton X-100, against collagen IV (microvascular marker) and amyloid-$\beta_{(1\text{-}40)}$ (1:100, rabbit polyclonal, ThermoFisher catalog# 44-348A) overnight at room temperature. Sections were washed 3x with PBS, then incubated with secondary antibodies conjugated to fluorophores for 2 hours at room temperature in the dark. Brain slices were then washed with PBS and mounted using Prolong Diamond antifade with DAPI (Thermofisher, catalog# P36971). Z-stacks were obtained using 40x water immersion objective (numerical aperture 0.9) on a Leica DM6 microscope coupled to a Crest X-Light V2 (CrestOptics, Rome, Italy) spinning-disk confocal microscope, attached to a high-sensitivity Evolve Delta EMCDD camera (Teledyne Photometrics, Tucson, AZ) controlled by uManager software in 512 x 512 pixels/fields of view at a pixel resolution of 0.290 nm (37). A total of 15 fields of view from the cortex were captured. To analyze co-localization (overlapping pixels at each channel or those in close proximity) between collagen IV and amyloid-β plaques, images were segmented and the area of overlap between channels were calculated using ImageJ's ImageCalculator plugin. Total area of overlap were subtracted from the total area of collagen IV labeling to provide a percentage of microvasculature co-localizing with amyloid-β, which we named Microvascular CAA (%).

The same protocol was followed to label pial artery CAA, except for the antibody to label pial arteries, which was an anti-smooth muscle myosin heavy chain 1 (1:200, rabbit polyclonal, Abcam, Cambridge, UK, catalog# ab124679).

**Pressure myography**

*5x-FAD* and WT littermates were euthanized with 4% inhalational isoflurane, followed by exsanguination and decapitation. The brain was excised and kept in an ice-cold tissue collection physiologic salt solution (PSS, in mmol/L): 140 NaCl, 5 KCl, 2 $MgCl_2$, 10 Dextrose, 10 HEPES, pH 7.4. The tissue collection PSS was supplemented with 0.05% bovine serum albumin (BSA, catalog # BP1600-100, Fisher Scientific, Waltham, MA). Pial arteries, including posterior communicating arteries, anterior communicating arteries or small branches from the middle cerebral arteries, were carefully dissected in a water-jacketed dissection dish connected to a circulating cold-water bath (4°C) filled with PSS. After isolation, arteries were mounted in a custom-made pressure myograph chamber filled with PSS[2]. The cannula was filled with vessel PSS (in mmol/L): 124 NaCl, 3 KCl, 2 $MgCl_2$, 1.085 $NaH_2PO_4 \bullet 2H_20$, 26 $NaHCO_3$, 1.8 $CaCl_2$, 4 Dextrose. Both ends were cannulated onto glass cannulas (30–50 µm diameter at the tip), one side was pressurized and the other was closed to generate a blind sac. The solution was constantly oxygenated with 21% $O_2$ / 5% $CO_2$ / balance $N_2$ to maintain pH 7.4. For endothelium-denudation experiments, an air bubble was slowly passed through the intraluminal side to damage the endothelial cell layer. The pressure myography chamber was transferred to a microscope and connected to a pressure-servo pump to control intraluminal pressure (Living Systems Instrumentation, Burlington, VT). The preparation was equilibrated for 20 minutes in warmed (37° C), oxygenated vessel PSS exchanged at a rate of 3–5 mL per minute at a pressure of 15 mmHg. Intraluminal pressure was then increased to 50 mmHg and equilibrated until the generation of spontaneous myogenic tone. Real-time data were recorded at 15 Hz using the IonWizard v7.3 software (IonOptix, Westwood, MA).

*Assessment of Pial arteries function*

Myogenic tone was calculated as: Myogenic tone (%) = [1 – (LDT/LDP)]*100, where LDT is the lumen diameter after the generation of spontaneous myogenic tone and LDP is the passive lumen diameter (after incubation in $Ca^{2+}$-free PSS). Preparations were then challenged with different compounds, no more than 1 compound per vessel. Iberiotoxin (IbTox, 30 nM, Anaspec, Fremont, CA, catalog# AS-60763) was used to evaluate the participation of $BK_{Ca}$ channels in myogenic tone regulation. Endothelin-1 (ET-1, 30nM, agonist of endothelin receptors), to evaluate G protein-coupled receptor-dependent vasoconstriction; 60 mM $[K^+]_{extracellular}$ (balanced NaCl) to evaluate receptor-independent depolarization and constriction. In a subset of experiments (Figure 4), the broad-spectrum reducing agent 1, 4-dithiothreitol (DTT, 10 μM) was used to rescue thiol motifs from oxidation. L-NAME (200 μM) was added to another set of pial arteries to assess the contribution of NOS isoforms to basal myogenic tone. All preparations were exposed to 60 mM $[K^+]_{extracellular}$ PSS at the end of the experiment to ensure viability after all treatments. After experimental protocols, the passive diameter was obtained by bathing all preparations in $Ca^{2+}$-free PSS (in mmol/L): 124 NaCl, 3 KCl, 2 $MgCl_2$, 1.085 $NaH_2PO_4$•$2H_20$, 26 $NaHCO_3$, 4 Dextrose, 2 EGTA, 0.01 diltiazem, 0.1 sodium nitroprusside.

A subset of pressurized arteries was subjected to a myogenic reactivity curve to assess autoregulation. After stabilization and development of spontaneous myogenic tone, intraluminal pressure was increased from 5 mmHg up to 160 mmHg, in 20 mmHg steps. Arteries were allowed to equilibrate at each pressure for 5 minutes before pressure was raised. At the end of the curve, arteries were exposed to 50 mmHg and bathed in $Ca^{2+}$-free PSS to collect passive diameter at each pressure step. Passive diameters were also used to assess structural and biomechanical parameters of pressurized pial arteries, as described previously[3, 4].

### Smooth Muscle Cell Isolation

Pial artery smooth muscle cells were isolated from *5x-FAD* and WT littermates for patch-clamp electrophysiology experiments. Pial arteries were dissected in ice-cold tissue PSS as described above, then transferred to vials containing 1 ml of tissue collection PSS (without BSA) supplemented with papain (1 mg/mL) and DTT (1 mg/mL) for 15 minutes at 37°C. After three washes with tissue PSS, the solution was exchanged by fresh tissue collection PSS supplemented with collagenase type II (1mg/mL) and again incubated for 15 minutes at 37˚C. Following three washes with Tissue PSS, arteries were gently triturated using a small diameter fire-polished glass Pasteur pipette.

### Recording of single channel $BK_{Ca}$ currents

Single channel $BK_{Ca}$ currents were recorded from freshly isolated pial artery smooth muscle cells in the inside-out excised patch configuration. Currents were acquired using a MultiClamp 700B (Molecular Devices, Sunnyvale, CA) amplifier at 10 kHz with a low-pass filter of 0.2 kHz. Single channel current data was acquired using DigiData 1550B (Molecular Devices) data acquisition system and pClamp 10.7 software (Axon Instruments, Molecular Devices). Microelectrodes pulled (P-1000, Sutter Instruments, Novato, CA) from 1.2mm x 0.94mm borosilicate glass capillary tubes (BF120-94-10, Sutter Instruments) were filled with a pipette solution containing (in mM) 140 KCl, 1 EDTA, 10 HEPES with pH adjusted to 7.3 with Tris. Bath solution consisted of (in mM) 140 KCl, 1 EDTA, 10 HEPES with pH adjusted to 7.3 with Tris and supplemented with $CaCl_2$ to achieve a 10 μM free $Ca^{2+}$ concentration. Single channel $BK_{Ca}$ currents were recorded for a duration of 30 seconds at each voltage step (+20, +40, +60 mV). $BK_{Ca}$ open probability (Po) data was analyzed using single-channel event analysis P(open) algorithm in Clampfit 10 (Axon Instruments, Molecular Devices). To evaluate whether 1, 4-dithiothreitol (DTT), a reducing agent, restored channel activity, $BK_{Ca}$ currents were also recorded in presence of vehicle or DTT (10 μM). The Po data were normalized by the total number of $BK_{Ca}$ present in the membrane patch when voltage was raised to +80 mV[5].

**Quantitative (q) PCR**

Messenger RNA expression of $BK_{\alpha}$ and $BK_{\beta 1}$, endothelial nitric oxide synthase (eNOS), neuronal NOS (nNOS), and inducible NOS (iNOS) were quantified from pial arteries and cortex from WT and *5x-FAD* mice using Taqman probes (**Table S1**). Briefly, the tissue was collected and stored in RNA Later (QIAGEN, Redwood City, CA, catalog# 76104) at -80°C until mRNA isolation. mRNA was extracted using RNeasy Plus Micro Kit (QIAGEN, catalog# 74034) following the manufacturer's instructions. mRNA to cDNA conversion was performed using iScriptTM Select cDNA Synthesis Kit (Bio-Rad, Hercules, CA, catalog# 170-8897), and 50 ng of cDNA were loaded into separate wells for qPCR amplification using TaqMan Fast Advanced Master Mix (Applied Biosystems, catalog# 4444557) in an Applied Biosystems 7300 Real-Time PCR System (Applied Biosystems, Foster City, CA). Samples were run in duplicates, and β-actin (actin) was used as endogenous control. Negative controls (RNA-free water) were included in all experiments. Cycling conditions consisted of an initial step of 50°C for 2 minutes and 10 minutes at 95°C followed by 50 cycles of 95°C for 15 seconds and 60°C for 1 minute. Gene expression is represented by the $2^{-\Delta\Delta CT}$.

**Western blot of pial arteries**

Pial arteries from *5x-FAD* and wild-type littermaters were carefully isolated, cleaned of pial meninges and homogenized in 80 µL of lysis buffer (50 mmol/L Tris-HCl (pH 7.4) containing 1% Nonited P-40, 0.5% sodium deoxycholate, 150 mmol/L NaCl, 1 mmol/L EDTA, 0.1% SDS, 1 mmol/L phenylmethylsulfonyl fluoride (PMSF), 1 µg/mL pepstatin A, 1 µg/mL leupeptin and 1µg/mL aprotinin). Lysates were then centrifuged at 4°C to collect the supernatant. Protein concentration in the lysate was measured using a Pierce™ BCA Protein Assay Kit (Thermo Fisher Scientific, catalog# 23225). A total of 20 µg of proteins were loaded into 10% SDS-PAGE and transferred to a nitrocellulose membrane. Membranes were blocked with Tris-buffered saline (TBS) containing 2% albumin and 0.01% Tween for 1 hour at room temperature. Membranes were then incubated with the anti-$BK_{\alpha}$ primary antibody (Alomone labs, Table S2) or anti-β-actin (Invitrogen, Table S2) overnight at 4°C. Membranes were washed with TBS and incubated with the respectively secondary antibodies conjugated with fluorescent probes for 1h at room temperature (Table S2). The fluorescent signal was recorded using the ChemiDoc MP Imaging System (Bio-Rad Laboratories Inc., Hercules, CA). Data are shown as a ratio of $BK_{\alpha}$ protein levels normalized to β-actin.

**Assessment of SMC $Ca^{2+}$ sparks**

*Reverse en-face cerebral artery preparation and time-lapse imaging of $Ca^{2+}$ sparks*

Mice were anesthetized via isoflurane and subsequently euthanized by decapitation to preserve the integrity of the intended blood vessels. Basilar and posterior cerebral arteries from *5x-FAD* and WT mice were prepared *reverse en face*, where the vessel is cut longitudinally with fine spring scissors and pinned onto a Sylgard block using insect pins (Austerlitz Insect Pins®, minutiens pin 0.10 mm in stainless steel) with the smooth muscle cells facing up. Vessels were loaded with the calcium indicator Cal-520 AM (10 µM, Abcam, catalog number# ab171868) with equi-volume 20% pluronic acid in tissue collection PSS for 30 minutes at 37°C. After loading, preparations were moved into a high-speed, high-resolution spinning-disk confocal microscope for time-lapse imaging of $Ca^{2+}$ sparks.

Reverse *en face* arterial preparations were utilized for time-lapse imaging of smooth muscle cell $Ca^{2+}$ sparks. Arteries were continuously supplied with warm (37°C) imaging PSS consisting of (in mmol/L): 2 ascorbic acid, 2.5 $CaCl_2$, 119 NaCl, 4.7 KCl, 0.5 $MgSO_4$, 1.18 $KH_2PO_4$,

10 Dextrose, 21 $NaHCO_3$, 10 HEPES, pH 7.4 via perfusion pump. The imaging dishes were fixed onto a Leica DM6 microscope with a 63x water immersion objective (numerical aperture 0.9). Time-lapse images were obtained via a high-sensitivity Evolve Delta EMCDD camera (Teledyne Photometrics, Tucson, AZ) controlled by µManager software in 512 x 512 pixels/fields of view at a pixel resolution of 0.290 nm[6]. Time-lapse images (1000 images) from each field of view were imaged for ~20 seconds to visualize intracellular $Ca^{2+}$ transients in smooth muscle cells. The frame rate was approximately 25 frames per second.

*Analysis of $Ca^{2+}$ Sparks*

Greyscale time-lapse imaging files were opened on custom-made software, kindly provided by Dr. Mark Nelson and Dr. Adrian Bonev from the University of Vermont (SparkAn version 5.5.6.0)[7]. Pixel-by-pixel fluorescence intensity of the first 10 images were averaged to generate background fluorescence ($F_0$). Individual smooth muscle cell $Ca^{2+}$ sparks were identified as described previously[8]: duration less than 0.7s, an amplitude greater than 1.1 ($\Delta F/F_0$) and a spatial spread area less than 25 µm$^2$. Then, theses parameters were quantified: event frequency (total events/ time) and the number of active sites per cell. A total of 5 regions of interest per preparations were randomly chosen by an investigator blinded to the experimental group, and the $Ca^{2+}$ spark frequency and sites per cell within the ROI were assessed.

**Oxidized Glutathione levels**

Global oxidative stress in brain lysates was assessed using a commercially available Kit (Glutathione colorimetric detection kit, catalog# K006-H1, Arbor Assays, Ann Arbor, MI) following the manufacturer's instructions. The colorimetric kit detects total glutathione (GSH) and oxidized glutathione (glutathione disulfide, GSSG) levels. Final expression values were further normalized by total concentration of proteins in the lysate to ensure that any observed were not due to higher protein abundance.

**iNOS and $BK_\alpha$ immunofluorescence**

To verify iNOS and $BK_\alpha$ localization in pial arteries and perform a semi-quantitative assessment of their expression, brain sections (50 µm) from perfusion-fixed brains were processed for immunolabeling as described above. Brain sections were incubated overnight with the primary anti-iNOS or anti-collagen IV antibodies (Table S2). Then, the sections were washed with PBS and incubated with specific secondary antibodies tagged with fluorescent probes diluted in 2% horse serum and 0.1% Triton X-100 in PBS (Table S2) for 2 hours at room temperature. Images were acquired using a Zeiss LSM880 laser scanning confocal microscope (Carl Zeiss Jena GmBH, Jena, Germany), using a 40x oil immersion objective (numerical aperture 1.3) to acquire Z-stacks of pial vessels positively labeled for collagen IV. Images were collected by an investigator blinded to experimental groups, and a total of 5 pial arteries randomly selected were imaged from each individual mouse. Images were processed using ImageJ to generate maximum-intensity projections. All brain sections were immunolabeled simultaneously to reduce the likelihood of potential artifacts. No post-acquisition processing was performed in the images, other than generation of maximum intensity projection maps, and settings for laser power, PMT gain, and offset were adjusted in one brain slice from negative control (no primary antibodies) and kept constant to image the remaining *5x-FAD*, WT, and no-labeling controls (negative control).

**Assessment of total S-Nitrosylated (S-NO) proteins**

Expression of S-nitrosylated proteins in brains from WT and *5x-FAD* were quantified using a "biotin-switch" method in the S-nitrosylated protein detection Kit (Cayman Chemical, Ann Arbor, MI, catalog# 10006518). Briefly, 25 mg of cortex brain tissue was homogenized in 500 µL of Buffer A containing blocking reagent and protease inhibitors for 30 minutes on ice to lyse cells and block the free thiol residues. The homogenate was then centrifuged for 10 minutes at 4ºC 15,000 x g. The supernatant was split into two aliquots, and the protein was precipitated by adding acetone, followed by incubation at -20ºC for 1 hour. After the incubation samples were centrifuged at 3,000 x g for 10 minutes at 4ºC, acetone was removed, and the pellet from one of the aliquots was re-suspended in buffer B alone to verify the contribution of signal from endogenously biotinylated protein (internal control aliquot). The other aliquot was re-suspended in Buffer B containing reducing and labeling reagents. This buffer reduces the S-nitrosothiols to yield free thiol(s), which were covalently labeled with maleimide-biotin. After an incubation period of 1h at room temperature, proteins were precipitated with acetone and used for Western blot or affinity-purification assays, as described below.

*Western blot*

Samples were centrifuged to remove acetone, and the protein pellet re-suspended in 150 μL of lysis buffer (50 mmol/L Tris-HCl (pH 7.4) containing 1% Nonited P-40, 0.5% sodium deoxycholate, 150 mmol/L NaCl, 1 mmol/L EDTA, 0.1% SDS, 1 mmol/L phenylmethylsulfonyl fluoride (PMSF), 1 μg/mL pepstatin A, 1 μg/mL leupeptin and 1μg/mL aprotinin). 20 μL were used to measure protein concentration using the BCA method (Pierce™ BCA Protein Assay Kit, Thermo Fisher Scientific, catalog# 23225). A total of 30 μg of proteins were separated on 10% SDS-PAGE and transferred to a nitrocellulose membrane. Membranes were blocked with Tris-buffered saline (TBS) containing 2% albumin and 0.01% Tween for 1 hour at room temperature. Membranes were incubated with the S-Nitrosylation Detection Reagent I for 1 hour, and the chemiluminescent signal was recorded using the ChemiDoc MP Imaging System (Bio-Rad Laboratories Inc., Hercules, CA). Membrane was then incubated with the primary antibody anti-β-actin (endogenous control, Table S2) overnight at 4°C. Membranes were washed 3x with TBS-Tween and incubated with specific secondary fluorescence-coupled antibodies (Table 2) for 1 hour at room temperature. Membranes were developed using ChemiDoc MP Imaging System. The results were expressed by S-Nitrosylated protein levels normalized to the β-actin.

*Affinity-purification assay*

Briefly, to evaluate $BK_{\alpha}$ S-NO, 500 μg of re-suspended proteins (after isolation of S-NO proteins as described above) were transferred to a 1.5 mL tube, and the total volume was adjusted to 200 μL. An aliquot of 20 μL from the original sample was saved to use as the input control sample. Then, 50 μL of streptavidin-coated magnetic beads (ThermoFisher Scientific, catalog# MSPB-6003-74) was added to the sample to bind to biotin-containing S-NO proteins and incubated for 4 hours at 4ºC in constant gentle agitation. After incubation, the tubes were moved to a magnetic base (DynaMag-2, catalog# 12321D, Invitrogen), followed by 3x washes with wash buffer. Then, proteins were eluted with 2x Laemmli buffer (4% SDS, 20% glycerol, 10% 2-mercaptoethanol, 0.004% bromphenol blue, and 0.125 M Tris HCl, pH 6.8), heated for 10 minutes at 100°C, loaded and separated by 10% SDS-PAGE, and proteins transferred to a nitrocellulose membrane. The membrane was incubated with a rabbit anti-$BK_{\alpha}$ antibody overnight at 4ºC and goat anti-β-actin. Membranes were washed 3x with Tris-buffered saline (TBS) containing 0.01 % Tween and incubated with specific secondary antibodies tagged with fluorescent probes (Table S2) for 1 hour at room temperature. Signals were detected using ChemiDoc MP Imaging System. Expression of $BK_{\alpha}$ was normalized by expression of β-actin.

*Co-localization of $BK_{\alpha}$ and S-NO*

Brain sections (50 µm) from perfusion-fixed mice were labeled with antibodies against BKα and the S-nitrosylation detection reagent conjugated with fluorescein. Briefly, sections were permeabilized, incubated with 10% horse serum (unspecific binding blocker) for 2 hours at room temperature, followed by overnight incubation with anti-$BK_{\alpha}$ antibody in 10% horse serum + 0.1% Triton X-100 at 4°C (Table 2). The next day, samples were washed buffer B containing 1% Triton X-100, reducing and labeling reagents for 1 hour at room temperature. Then, sections were washed and incubated with the S-nitrosylation detection reagent conjugated with fluorescein. Brain sections were then washed with Buffer B containing 1% Triton X-100, counterstained with DAPI (nuclei stain, Sigma-Aldrich, catalog# D9542), and mounted on glass slides using Fluoroshield mounting media. Slides were imaged using a Zeiss LSM880 laser scanning confocal microscope (Carl Zeiss Jena GmBH, Jena, Germany), using a 40x oil immersion objective (numerical aperture 1.3) to acquire Z-stacks of pial vessels. Images were collected by an investigator blinded to experimental groups, and a total of 5 pial arteries randomly selected were imaged from each individual mouse. Images were processed using ImageJ to generate maximum-intensity projections. All brain sections were immunolabeled simultaneously to reduce the likelihood of potential artifacts. No post-acquisition processing was performed in the images, other than generation of maximum intensity projection maps, and settings for laser power, PMT gain, and offset were adjusted in one brain slice from negative control (no primary antibodies) and kept constant to image the remaining *5x-FAD*, WT, and no-labeling controls (negative control). The vessel area was delimitated, and the results were expressed as the integrated density of fluorescence divided by the delimitated area of interest.

Co-localization was assessed as described previously by Dunn et al[9]. Briefly, each Z slice was considered separately in the Otsu threshold image, no- maximum-intensity projections were generated. Then, the green and red pixels intensity, Pearson's coefficient, M1 and M2 Manders' coefficient, and fluorogram were calculated as described previously[9]. Imaging of pial arteries was performed in a blinded and randomized manner, a total of 5 arteries were imaged per mouse for a total of 5 mice per group.

**_Post-mortem_ human brain samples**

*Post-mortem* human brain tissue were kindly provided from participants in the Banner Sun Health Research Institute Brain and Body Donation Program Sun City, Arizona[10]. All samples were collected within a short *post-mortem* interval (approximately 3h) and stored following gold-standard procedures to ensure maximal tissue viability for molecular / biochemical assessments (for details, please see[10-12]). For the purposes of this study, twenty-four (24) age-matched cortex brain samples were provided, 12 from non-AD participants (6 men and 6 women) and 12 diagnosed with AD (6 men and 6 women). In the non-AD group, we observed that a total of 6 participants (2 men and 4 women) had cerebral amyloid angiopathy (CAA), thus we excluded them from the non-AD group and created a third group: CAA. The age average of the groups are: non-AD: 81 ± 6.75 years; non-AD with CAA: 82.4 ± 6.87 years and AD: 84.6 ± 4.75 years. At the day of experiment, 50 mg of brain tissue were weighted and lysed, and supernatants were used to detected $BK_\alpha$ expression by Western blot and $BK_\alpha$ S-NO by affinity-purification followed by Western blot, following the protocols described previously for mouse samples.

**Cerebral hemodynamics by laser speckle contrast imaging**

*Basal cortical perfusion*

Mice were anesthetized with 3% isoflurane mixed with breathing air. After deep anesthesia, mice were intubated *via* tracheotomy for constant mechanical ventilation (RoVent Small Animal Ventilator, Kent Scientific Corporation, Torrington, CT). Mouse was then moved to a self-heated stereotaxic frame (Stoelting Co., Wood Dale, IL) to ensure head immobilization using ear bars. Body temperature was maintained constant at 37 ± 1°C using a rectal probe that provided feedback to the heating element of the stereotaxic frame. The scalp was shaved and exposed, a midline incision was performed to expose the skull, and the periosteum was gently removed with cotton-tipped applicators. The laser speckle contrast imaging system (PSI-Z; Perimed AB, Järfälla, Sweden) was placed 11-12 cm above the skull, and basal cerebral perfusion was recorded for 3 min at a rate of 20 images/s. All perfusion data were acquired using the manufacturer's software (PIMsoft v. 1.6; Perimed) and are expressed as perfusion units (PU)[4].

Cortical perfusion was analyzed considering standardized regions of interest: frontal, parietal, zone of pial anastomoses (ZPA) and temporal cortices of both hemispheres, as described previously[13]. These regions were evaluated separately because each is supplied by a different major cerebral artery and has discrete areas of anastomoses and compensatory capacity[13, 14]. Briefly, after the frontal pole limit (FP), the bregma and lambda points on the skull were localized, and a middle line was drawn to connect the frontal pole limit to lambda, passing through the bregma. Then, an area delimited from 1.5 mm from this midline and by the frontal pole and bregma was considered the frontal cortex region. The area delimited from 1.5 mm from bregma and 2 mm from lambda was considered the parietal cortex. The area 1 mm from the delimitation of the frontal and parietal region was designated as the ZPA. The remaining lateral area corresponds to the temporal cortex for both hemispheres. For a diagram of all different areas, see Figure S12.

*Neurovascular coupling following whiskers stimulation*

Neurovascular reactivity, an index of neurovascular coupling in the brain, was assessed by real-time increases in perfusion atop the whisker barrel cortex during mechanical whisker stimulation, as previously described[4, 15]. Following basal cerebral perfusion measurements, the skull was thinned atop the left whisker barrel cortex[16]. Then, the isoflurane dosage was lowered to 1.5% to reduce the influence of anesthesia on the physiological cardiovascular function while providing an acceptable level of anesthesia[17], including observable hemodynamic responses after stimulation[18]. Basal cerebral perfusion was allowed to stabilize for 15 minutes, then the contralateral vibrissae were stimulated with a brush at a rate of 5 Hz for 30 s, followed by a 3 minutes recovery period. This procedure was repeated three times, followed by one ipsilateral vibrissae stimulation to certify that there were no movement artifacts during the recordings. The recordings were acquired in real-time using the PSI-Z system at a frequency of 20 frames/s and analyzed using the PIMsoft software. The baseline was considered as the average of the 10 seconds before the vibrissae stimulus. Functional hyperemia was assessed as the average perfusion during the entire duration of the stimulation (30 s). In a subset of WT mice, 100 µl of a warm (37°C) 30 nM Iberiotoxin solution (dissolved in aCSF), or vehicle (aCSF) was added atop the thinned-skull cranial window to acutely block $BK_{Ca}$. After a 5 minutes incubation, the contralateral vibrissae were again stimulated. The iberiotoxin effect on the neurovascular reactivity was evaluated by comparing the functional hyperemia response before and after iberiotoxin or vehicle. Data are shown as the percent increase in perfusion from baseline.

**Chemicals**

Chemical reagents used in this study were purchased from Sigma-Aldrich unless otherwise indicated.

**Statistical analyses**

Data are expressed as means ± SEM. All data analyses were performed with GraphPad Prism 10. Differences between the means of the two experimental groups were analyzed with two-tailed *Student's* t-tests or Mann–Whitney test if the data did not follow a normal distribution. $BK_{Ca}$ Po data were analyzed by ordinary two-way ANOVA followed by a Sidak correction for multiple comparisons, or Mixed-Model ANOVA for paired experiments, followed by a Sidak correction for multiple comparisons. Human data in Figure 6 did not followed a Gaussian distribution, and were thus analyzed by a Brown-Forsythe and Welch ANOVA test with a Dunnett T3 correction for multiple comparisons. All tests used in the graphs are specified in each Figure legend. All data were tested for the possible presence of outliers by the ROUT method, which is based on a false discovery rate with a Q = 1. No outliers were detected in our data set. Absolute p values for all tests run, rather than symbols, are shown in Figures. Representative traces of perfusion shown in Figure 7 were smoothed in GraphPad using a 5 neighbors averaging method and a second polynomial transformation (less smooth).

**Table S1.** *Taqman probes used in this study.*

| *Target* | *ID Taqman assay* | *Catalog number* |
|---|---|---|
| BKα | Mm01268569_m1 | 4453320 |
| BKβ | Mm00466621_m1 | 4448892 |
| eNOS | Mm00435217_m1 | 4453320 |
| nNOS | Mm01208059_m1 | 4453320 |
| iNOS | Mm00440502_m1 | 4453320 |
| Actin | Mm00607939_s1 | 4453320 |

**Table S2.** *Antibodies used in this study.*

| ***Target*** | ***Manufacturer*** | ***Catalog number*** | ***dilution*** |
|---|---|---|---|
| Collagen IV | Novus Biologicals | NBP1-26549 | 1:200 |
| Amyloid-β$_{(1-40)}$ | ThermoFisher | 44-348A | 1:100 |
| BK$_{\alpha}$ | Alomone labs | APC-021 | 1:100-IF;<br>1:500 - WB |
| iNOS | Cell Signaling | #13120 | 1:100 |
| β-actin | Invitrogen | #PA1-183 | 1:5000 |
| α-smooth muscle actin | ThermoFisher | #PA5-18292 | 1:500 |
| donkey anti-rabbit AlexaFluor 488 | Jackson ImmunoResearch | # 711-545-152 | 1:800 |
| donkey anti-goat AlexaFluor 594 | Jackson ImmunoResearch | # 705-585-147 | 1:800 |
| donkey anti-rabbit AlexaFluor 594 | Jackson ImmunoResearch | # 711-585-152 | 1:800 |

**Supplemental Figures Legends**

**Figure S1.** *Sex-dependent amyloid-β accumulation in 5x-FAD.* **A-B)** Representative images (**A**) of thioflavin-T labeling of parenchymal "senile" amyloid-β plaques (green) in the cortex (left) and hippocampus (right) from female (top panels) and male (lower panels) *5x-FAD*. Quantification of those data show a trend towards increased amyloid-β accumulation in female *5x-FAD*, as shown in **B**. Bar = 100 µm. Statistical analysis: one-way ANOVA with a Sidak correction for multiple comparisons. **C-D)** Representative maximum intensity projection confocal images (**C**) showing immunolabeling for amyloid-β and the microvascular marker collagen IV (red) in the brains of females (top panels) and males (lower panels) *5x-FAD*. Bar = 30 µm. Note the low abundance of intraparenchymal CAA in the model, and no significant sex-differences were observed in microvascular CAA in *5x-FAD*, as evidenced in the summary graph (**D**). Each data point represents the average of 5 random fields of view within the cortex of an individual mouse. Statistical analysis: unpaired two-tailed Mann-Whitney test. **E-F)** Representative images of pial artery CAA (**E**) showing perivascular amyloid-β plaques (green) surrounding smooth muscle actin-positive arteries (α-SMA, red) in both females (top panels) and males (lower panels). Bar =

30 µm. No significant sex-differences were observed in pial artery CAA in *5x-FAD*, as shown in the summary graph (**F**). Each data point represents the average of all pial arteries found in the brain slice of an individual mouse. Statistical analysis: unpaired two-tailed Mann-Whitney test. All data are means ± SEM. Data in **F** are a re-analysis of original findings published by our laboratory[2].

**Figure S2.** *Autoregulation is maintained in pial arteries from 5x-FAD.* **A)** Myogenic reactivity, assessed as myogenic tone at each pressure step within the autoregulatory curve, was not different between female *5x-FAD* and WT littermates. N = 9 pial arteries per group, one artery per mouse. Statistical analyses: ordinary two-way ANOVA with a Sidak correction for multiple comparisons. **B)** Area under the curve for the data shown in **A**. Statistical analyses for areas under the curve: unpaired two-tailed *Student*'s t-test. **C)** Similarly, myogenic reactivity was not significantly different between male *5x-FAD* and WT littermates. N = 7 (WT) and 9 (*5x-FAD*) arteries, one artery per mouse. Statistical analyses: ordinary two-way ANOVA with a Sidak correction for multiple comparisons. **D)** Area under the curve for the data shown in **C**. Statistical analyses for areas under the curve: unpaired two-tailed *Student*'s t-test. Data are means ± SEM.

**Figure S3.** *No significant structural remodeling was observed in pial arteries of 5x-FAD.* **A-D)** No significant differences were observed in outer (**A**) and lumen (**B**) diameters, wall thickness (**C**) or wall-to-lumen ratio (**D**) of pial arteries isolated from female *5x-FAD* or WT littermates. N = 12 (WT) and 13 (*5x-FAD*) pial arteries per group, one artery per mouse. **E-H)** Similarly, no significant structural differences were observed in pial arteries isolated from male *5x-FAD* or WT littermates. N = 10 (WT) and 11 (*5x-FAD*) arteries, one artery per mouse. Statistical analyses: ordinary two-way ANOVA with a Sidak correction for multiple comparisons. Data are means ± SEM.

**Figure S4.** *Biomechanical properties of pial arteries are similar between 5x-FAD and WT littermates.* **A-D)** No significant differences were observed in distensibility (**A**), stress (**B**), vascular compliance (**C**, assessed by the stress-strain relationship) or vascular stiffness (**D**, quantified as the β-coefficient of individual stress-strain relationships). N = 12 (WT) and 13 (*5x-FAD*) pial arteries per group, one artery per mouse. (**E-H)** Similarly, biomechanical properties of pial arteries isolated from male *5x-FAD* or WT littermates were identical. N = 10 (WT) and 11 (*5x-FAD*) arteries, one artery per mouse. Statistical analyses for **A, B, E, F:** ordinary two-way ANOVA with a Sidak correction for multiple comparisons. Statistical analyses for **C, G**: three-way ANOVA. Statistical analyses for **D, H:** unpaired two-tailed Mann-Whitney t-test. All data are means ± SEM.

**Figure S5.** *mRNA expression of different $BK_{Ca}$ subunits.* **A-B)** Expression of $BK_{\beta 1}$ mRNA in pial artery lysates from female (**A**) and male (**B**) *5x-FAD* and WT littermates. No significant differences were observed in mRNA expression, although there was a trend towards a decrease in $BK_{\beta 1}$ mRNA in lysates from male *5x-FAD* when compared to WT littermates. Each data point is 1 pial artery lysate from an individual mouse. Statistical analysis: unpaired two-tailed *Student*'s t-test. **C-F)** Expression of $BK_{\alpha}$ and $BK_{\beta 1}$ subunits in cortical lysates from female (**C-D**) and male (**E-F**) *5x-FAD* and WT littermates. There was a significant lower expression of $BK_{\alpha}$ mRNA in cortical lysates of female *5x-FAD* when compared to WT littermates, and a trend towards an increase in $BK_{\beta 1}$ mRNA in male *5x-FAD*. Each data point is 1 cortical lysate from an individual mouse. Statistical analysis: unpaired two-tailed *Student*'s t-test. All data are means ± SEM.

**Figure S6.** *Western blot for $BK_{\alpha}$ protein expression in pial artery lysates.* Whole blot of the representative inserts shown in Figure 3B and 3D. Note the band for $BK_{\alpha}$ (top rectangle) and β-actin (lower rectangle). A total of 2 different blots were performed, all samples are from lysates obtained from individual mice. Data are means ± SEM.

**Figure S7.** *No evidence for oxidative stress or oxidative $BK_{Ca}$ PTM in male 5x-FAD.* **A-C)** Total glutathione (GSH, **A**), oxidized glutathione (GSSG, **B**) or the ratio between GSSG / total GSH (**C**) in brain lysates were not significantly between male *5x-FAD* and WT littermates. Each data point represents 1 brain lysate from an individual mouse. Statistical analyses: unpaired two-tailed *Student*'s t-test. Data are means ± SEM. **D-E)** Representative traces (**D**) and summary data (**E**) showing that incubation of pial artery smooth muscle cells with DTT (10 μM) does not recover single channel $BK_{Ca}$ Po in *5x-FAD*. Statistical analysis: matching mixed-model two-way ANOVA with a Šidák correction for multiple comparisons. N = 12 cells from 5 individual mice. Data are means ± SEM.

**Figure S8.** *mRNA expression of eNOS and nNOS in pial artery lysates from females.* **A)** Expression of endothelial NOS (eNOS) was not different in pial arteries from female *5x-FAD* and WT littermates. **B)** Similarly, expression of neuronal NOS (nNOS) was not different in pial arteries from *5x-FAD* when compared to WT littermates. Each data point represents 1 pial artery lysate from an individual mouse. Statistical analyses: unpaired two-tailed *Student*'s t-test. Data are means ± SEM.

**Figure S9.** *Global protein S-NO in cortical lysates of male mice.* **A)** Representative blots of β-actin (left) and global protein S-NO (right) from brain lysates of male *5x-FAD* and WT littermates. **B)** Summary data showing that global protein S-NO was significantly lower in male *5x-FAD* when compared to WT littermates. Each data point represents 1 lysate of the cortex of 1 mouse. Statistical analysis: unpaired two-tailed Mann-Whitney t-test. Data are means ± SEM>

**Figure S10.** *Whole blots of inserts shown in Figure 5.* **A-B)** Representative Western blot membrane of the inserts shown in Figure 5F. Samples underwent affinity-purification of S-NO proteins followed by Western blot. The rectangle in the top membrane (**A**) highlights the band for $BK_{\alpha}$, the rectangle in the lower membrane (**B**) shows the band for β-actin. Each lane is a cortical lysate from 1 individual mouse. The sample on the outer right lane was discarded due to artifacts during imaging of the membrane. Endogenous control: endogenously biotinylated proteins that did not undergo "biotin-switch".

**Figure S11.** *Whole blots of inserts shown in Figure 6.* **A)** Representative Western blot membrane of the insert shown in Figure 6A, probed for $BK_{\alpha}$ (top rectangle) and β-actin (lower rectangle). **B)** Representative Western blot membrane of the inserts shown in Figure 6C. Samples underwent affinity-purification (AP) of S-NO proteins followed by Western blot against BKα and β-actin. Each lane is a cortical lysate from 1 individual patient. Endogenous control: endogenously biotinylated proteins that did not undergo "biotin-switch".

**Figure S12.** *Basal cortical perfusion assessed by laser speckle contrast imaging.* **A)** Diagram of the regional cortical perfusion assessment showing the rectangles used to quantify perfusion in the different vascular territories. Lines were drawn using the greyscale image (left), then transferred to the pseudocolored perfusion maps (right). PC: parietal cortex; FC: frontal cortex; ZPA: zone of pial anastomoses; TC: temporal cortex; WC: whole cortex. “Created with BioRender.com”. **B-C)** Representative perfusion maps (**B**) and summary data (**C**) showing perfusion in the different cortical regions of female WT and 5x-FAD. Perfusion to the frontal cortex was significantly lower in female *5x-FAD* when compared to WT littermates, without differences in other regions. **D-E)** Representative perfusion maps (**D**) and summary data (**E**) showing perfusion in the different cortical regions of male WT and *5x-FAD*. No significant differences in perfusion were observed in male mice. Each data point in the summary graphs represents 1 mouse. Statistical analyses: ordinary two-way ANOVA with a Sidak post-hoc correction for multiple comparisons. Data are means ± SEM.

**Figure S13.** *Graphical abstract of the main findings of this study.* Microvascular and neurovascular impairments are present in both female and male *5x-FAD* when compared to WT littermates, albeit due to different molecular / cellular mechanisms. In females, we describe a partially reversible nitro-oxidative PTM of the $BK_{\alpha}$ subunit likely caused by excessive NO produced *via* iNOS in the presence of a pro-oxidative environment. On the other hand, in males, $BK_{Ca}$ impairment is a consequence of reduced expression of the $BK_{\alpha}$ subunit coupled to a lower frequency of $Ca^{2+}$ sparks, which are upstream and necessary for $BK_{Ca}$ gating. Created with BioRender.

**Movie S1.** *$Ca^{2+}$ sparks in reverse en face preparations of a pial artery from a female WT mouse*. Bar = 20 μm. The arrows indicate sites of $Ca^{2+}$ sparks. Pial arteries were loaded with Cal-520 AM (10 μM) and time-lapse recordings performed using high-speed, high-resolution spinning-disk confocal microscopy.

**Movie S2.** *$Ca^{2+}$ sparks in reverse en face preparations of a pial artery from a female 5x-FAD*. Bar = 20 μm. The arrows indicate sites of $Ca^{2+}$ sparks. Pial arteries were loaded with Cal-520 AM (10 μM) and time-lapse recordings performed using high-speed, high-resolution spinning-disk confocal microscopy.

**Movie S3.** *$Ca^{2+}$ sparks in reverse en face preparations of a pial artery from a male WT mouse*. Bar = 20 μm. The arrows indicate sites of $Ca^{2+}$ sparks. Pial arteries were loaded with Cal-520 AM (10 μM) and time-lapse recordings performed using high-speed, high-resolution spinning-disk confocal microscopy.

**Movie S4.** *$Ca^{2+}$ sparks in reverse en face preparations of a pial artery from a male 5x-FAD*. Bar = 20 μm. The arrows indicate sites of $Ca^{2+}$ sparks. Pial arteries were loaded with Cal-520 AM (10 μM) and time-lapse recordings performed using high-speed, high-resolution spinning-disk confocal microscopy.
