## Supplementary material for "Sex-specific mechanisms of cerebral microvascular BK_Ca_ dysfunction in a mouse model of Alzheimer’s disease": SI Figures 1 - 13

**Figure S1.** *Sex-dependent amyloid-β accumulation in 5x-FAD*

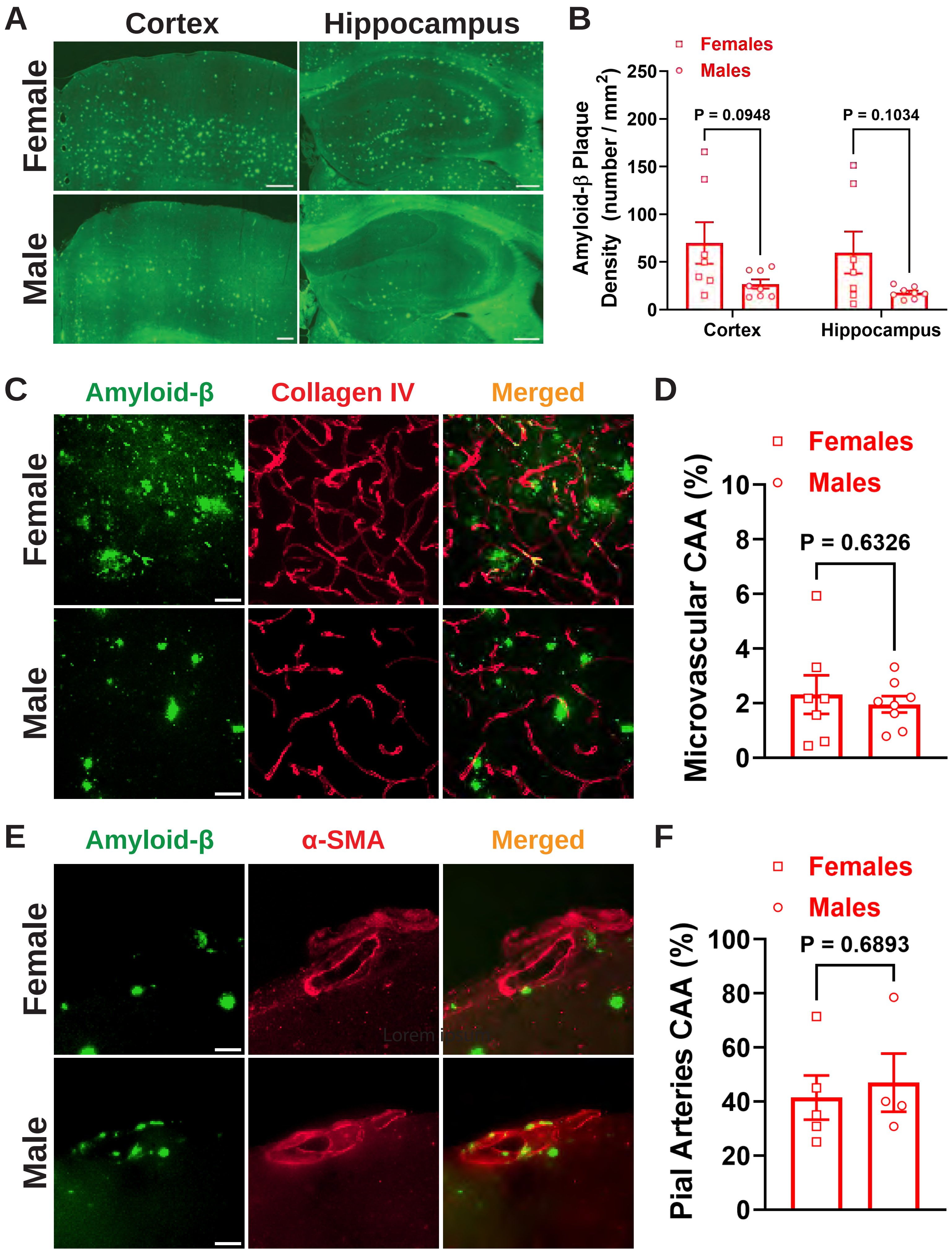

**Figure S2.** *Autoregulation is maintained in pial arteries from 5x-FAD*

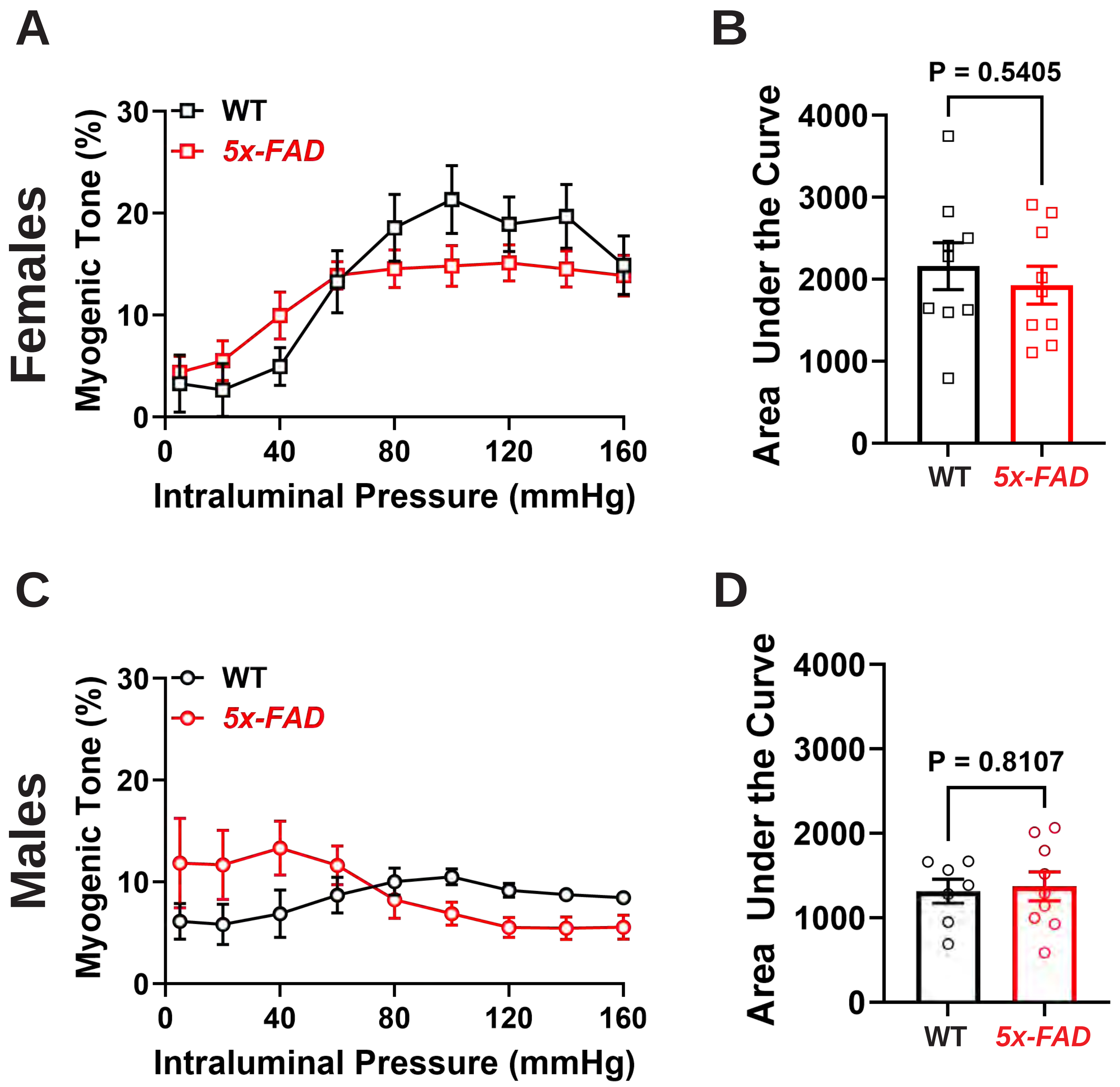

**Figure S3.** *No significant structural remodeling was observed in pial arteries of 5x-FAD*

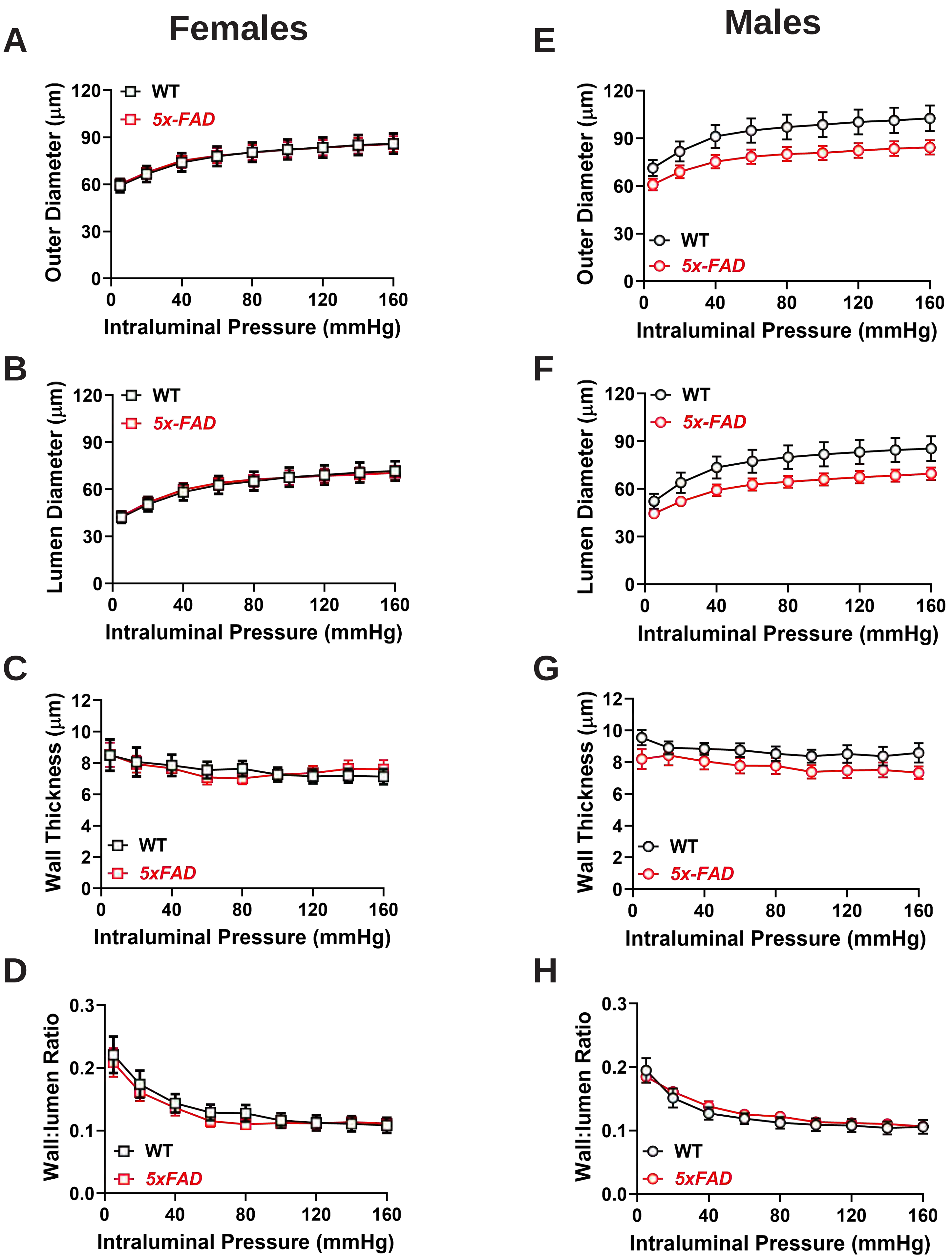

**Figure S4.** *Biomechanical properties of pial arteries are similar between 5x-FAD and WT littermates.*

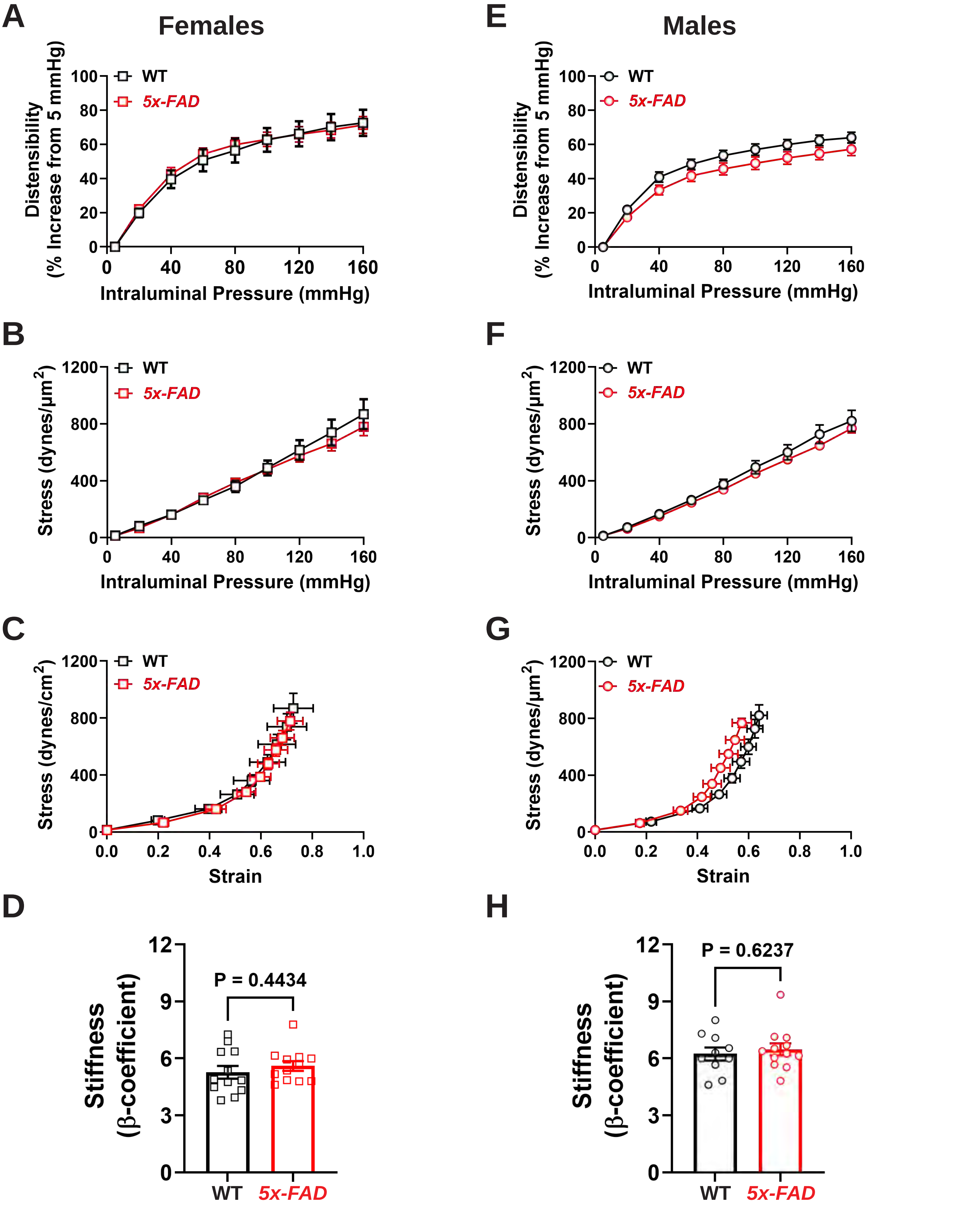

**Figure S5.** *NmRNA expression of different BKCa subunits*

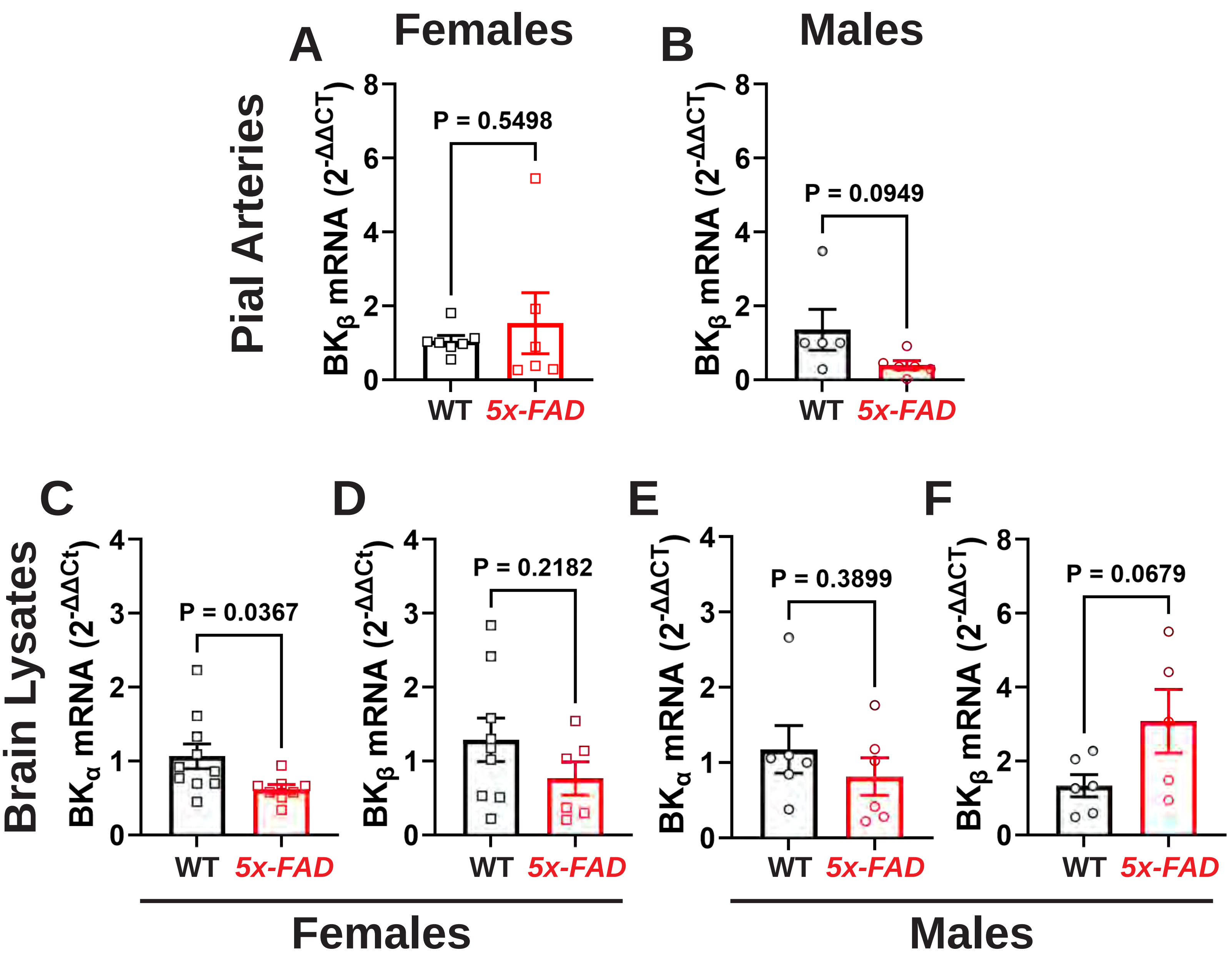

**Figure S6.** *Western blot for $BK_{\alpha}$ protein expression in pial artery lysates*

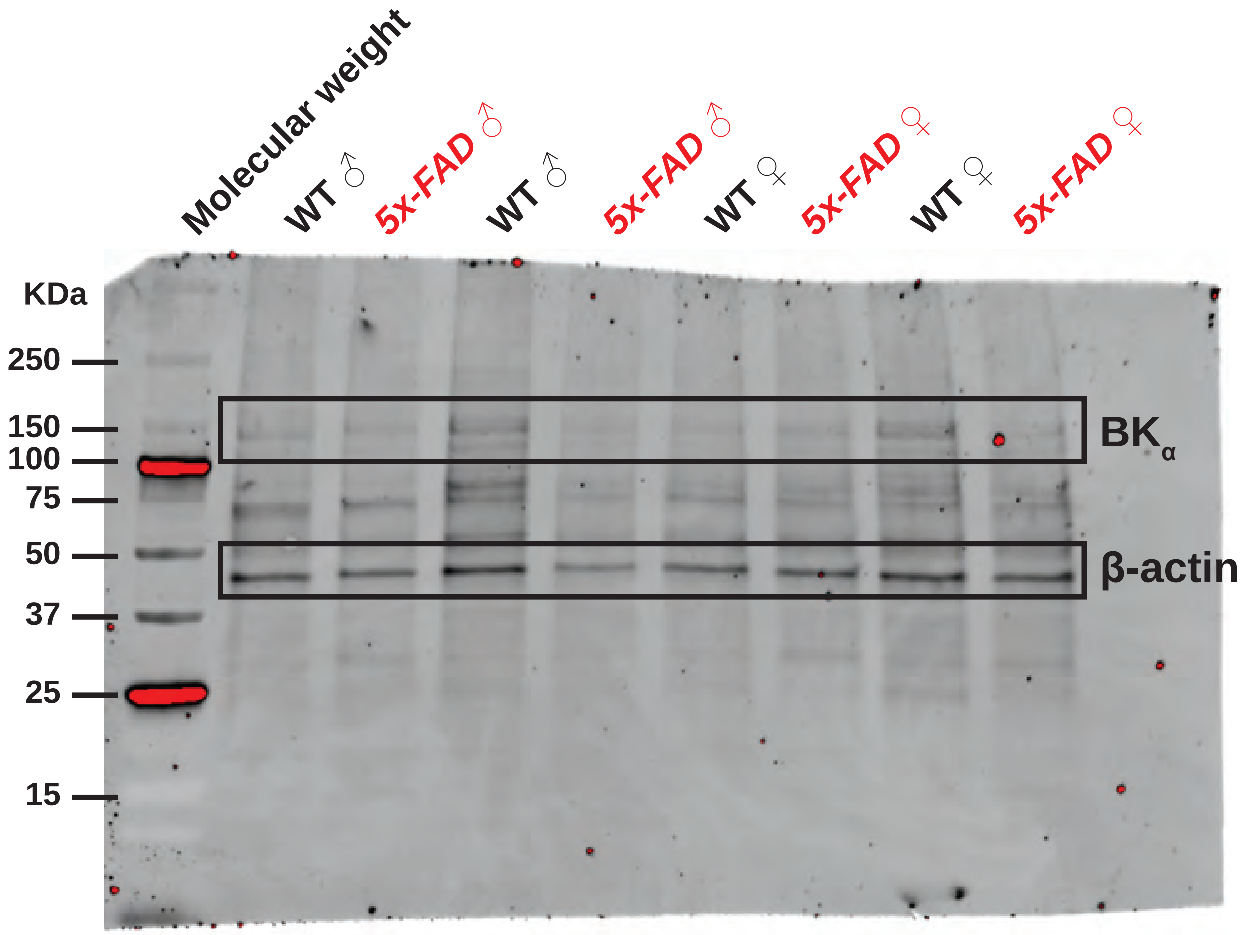

**Figure S7.** *No evidence for oxidative stress or oxidative $BK_{Ca}$ PTM in male 5x-FAD*

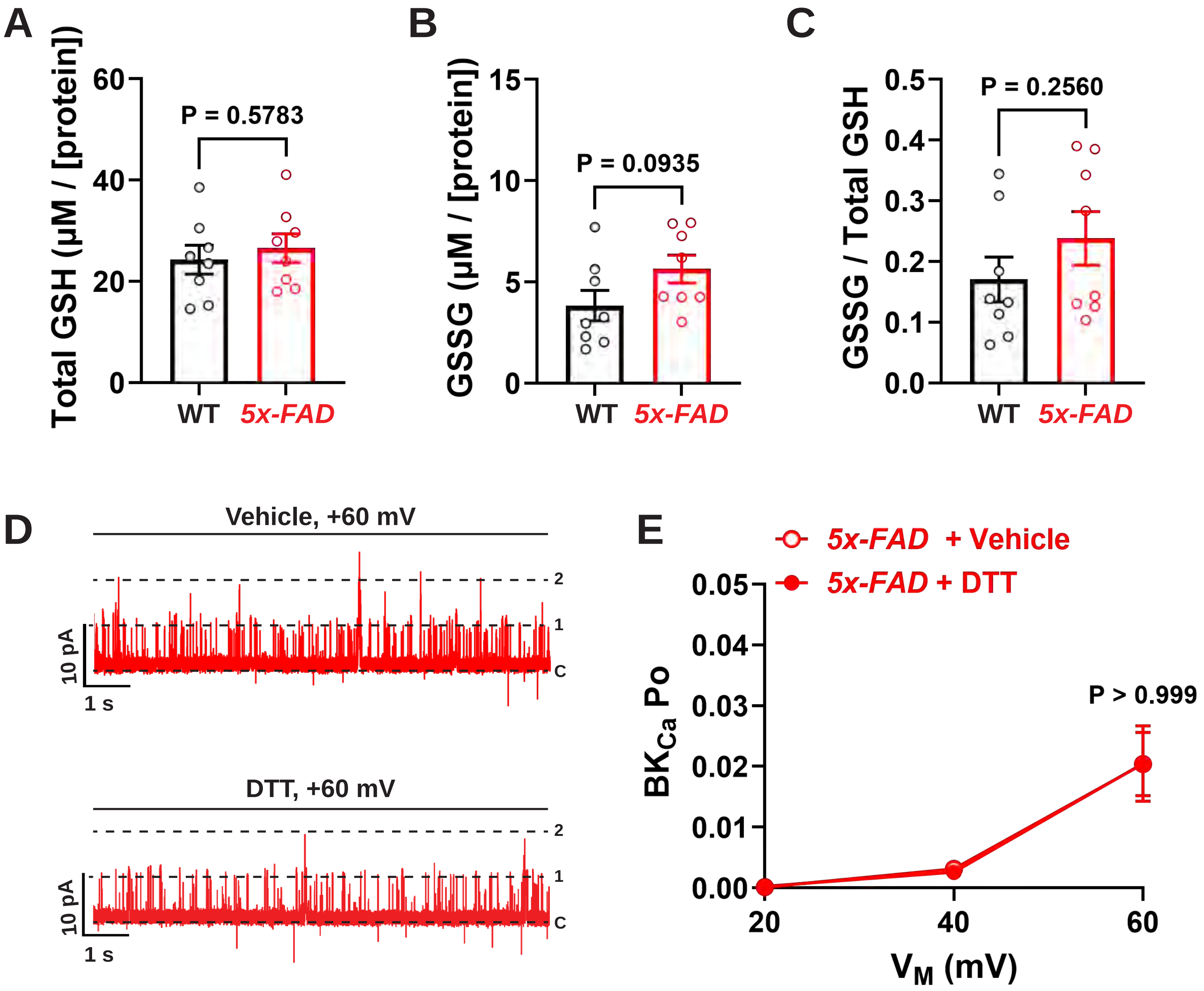

**Figure S8.** *mRNA expression of eNOS and nNOS in pial artery lysates from females*

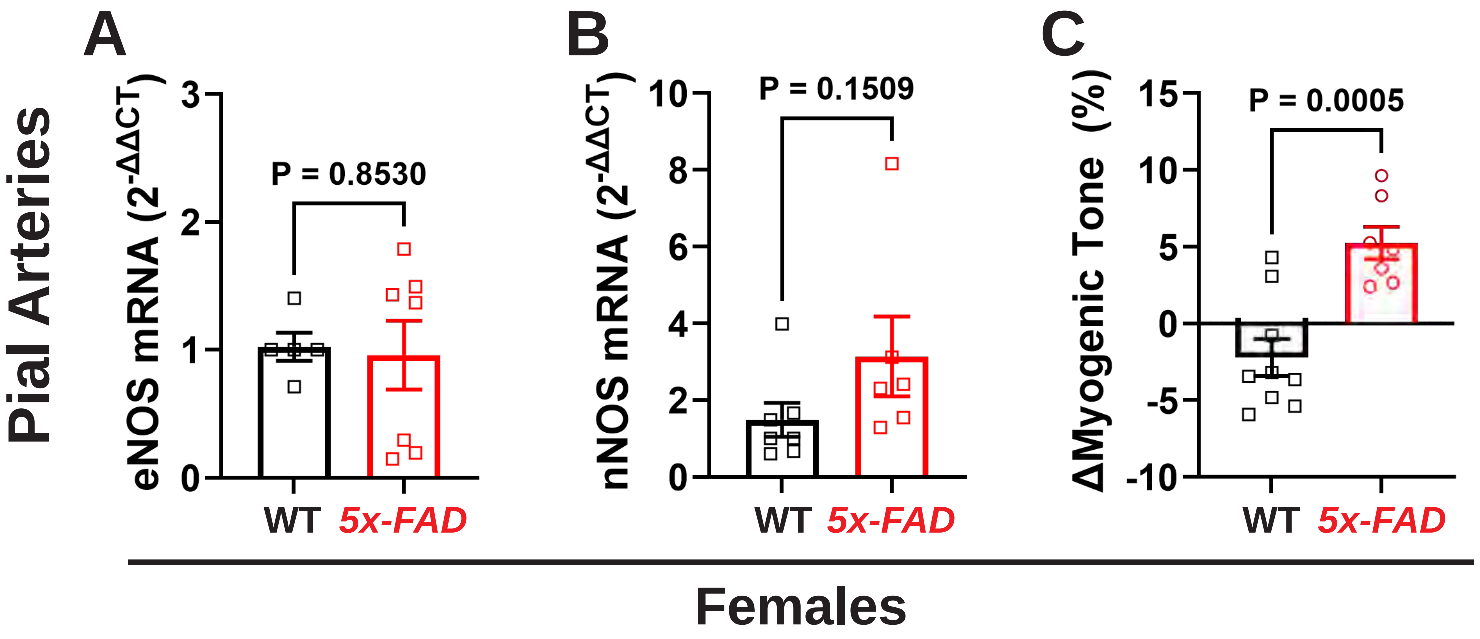

**Figure S9.** *Global protein S-NO in cortical lysates of male mice.*

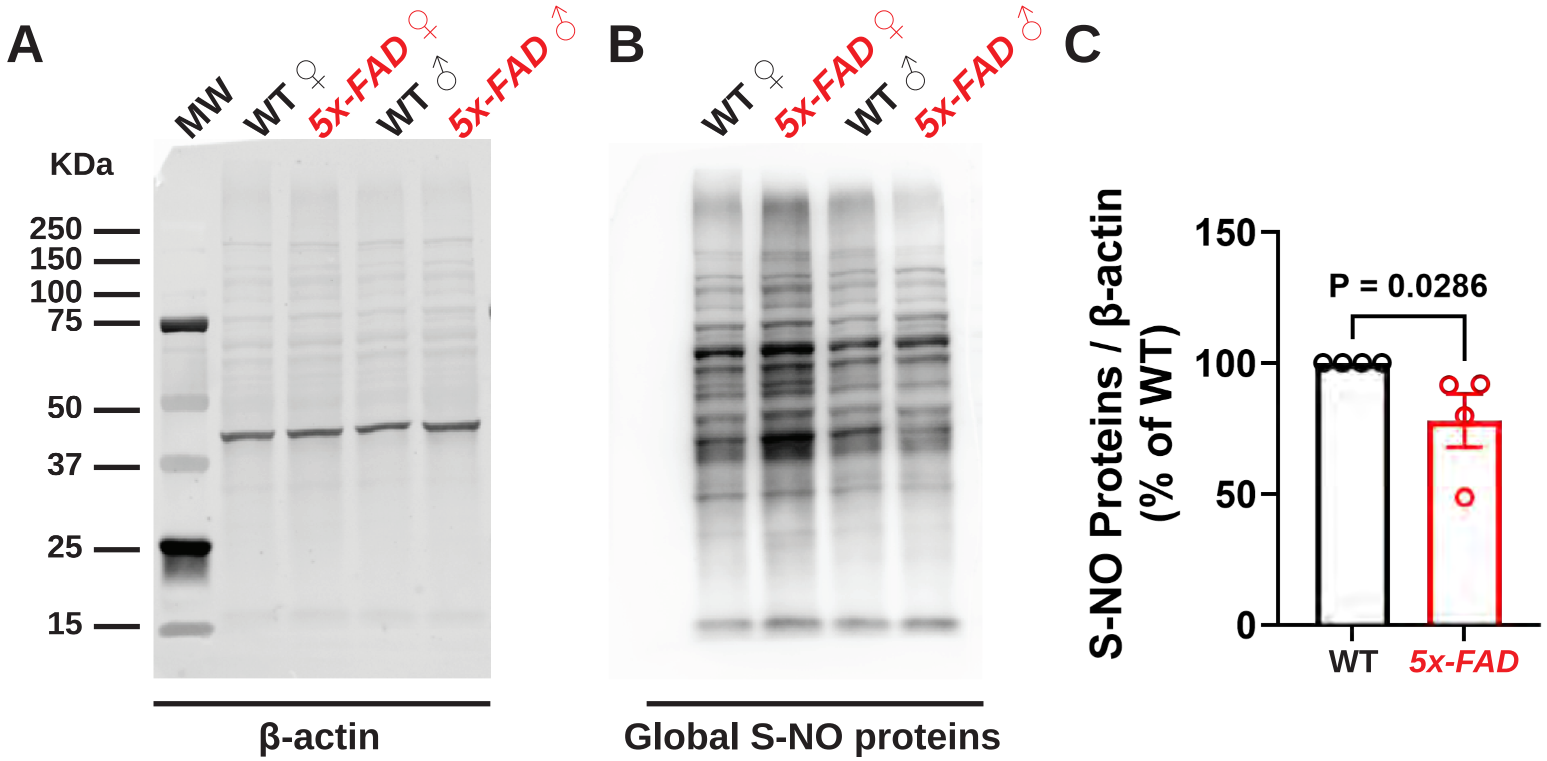

**Figure S10.** *Whole blots of inserts shown in Figure 5*

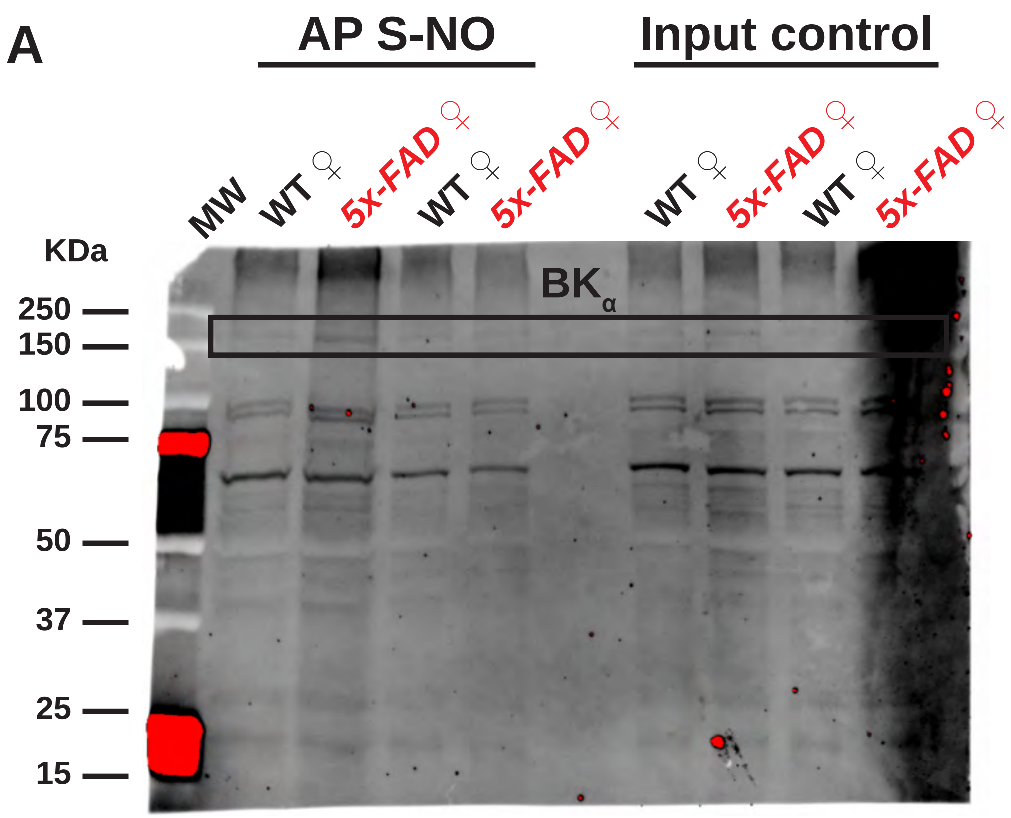


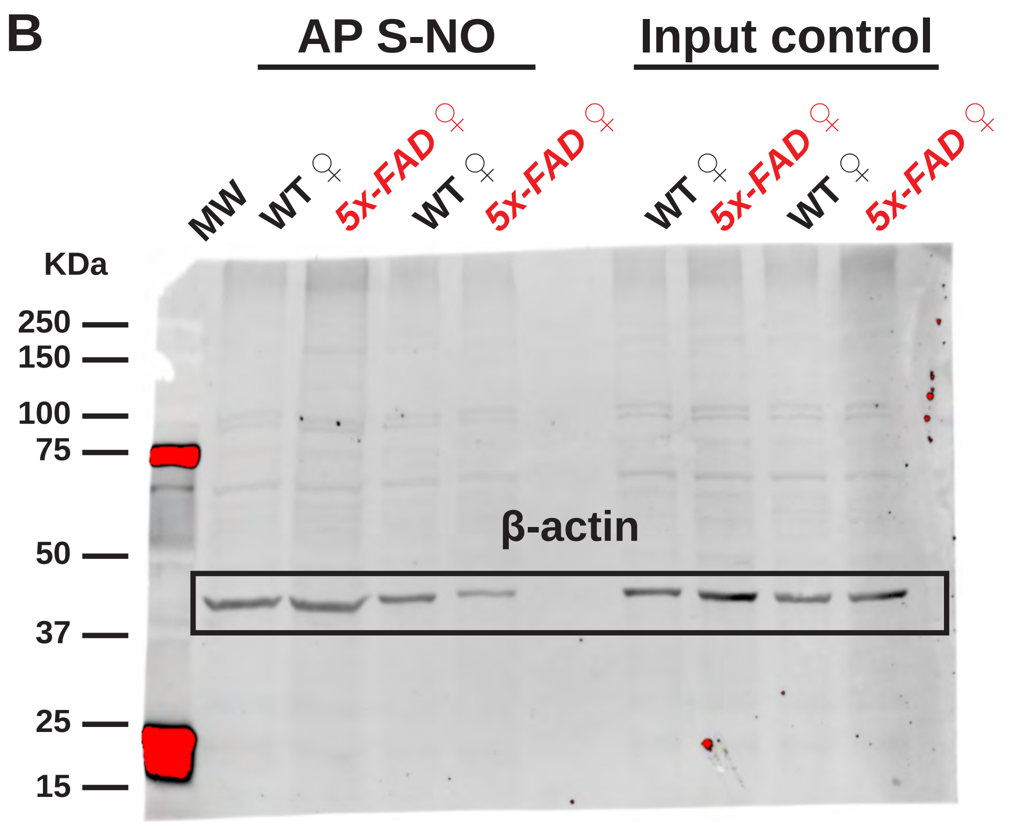

**Figure S11.** *Whole blots of inserts shown in Figure 6*

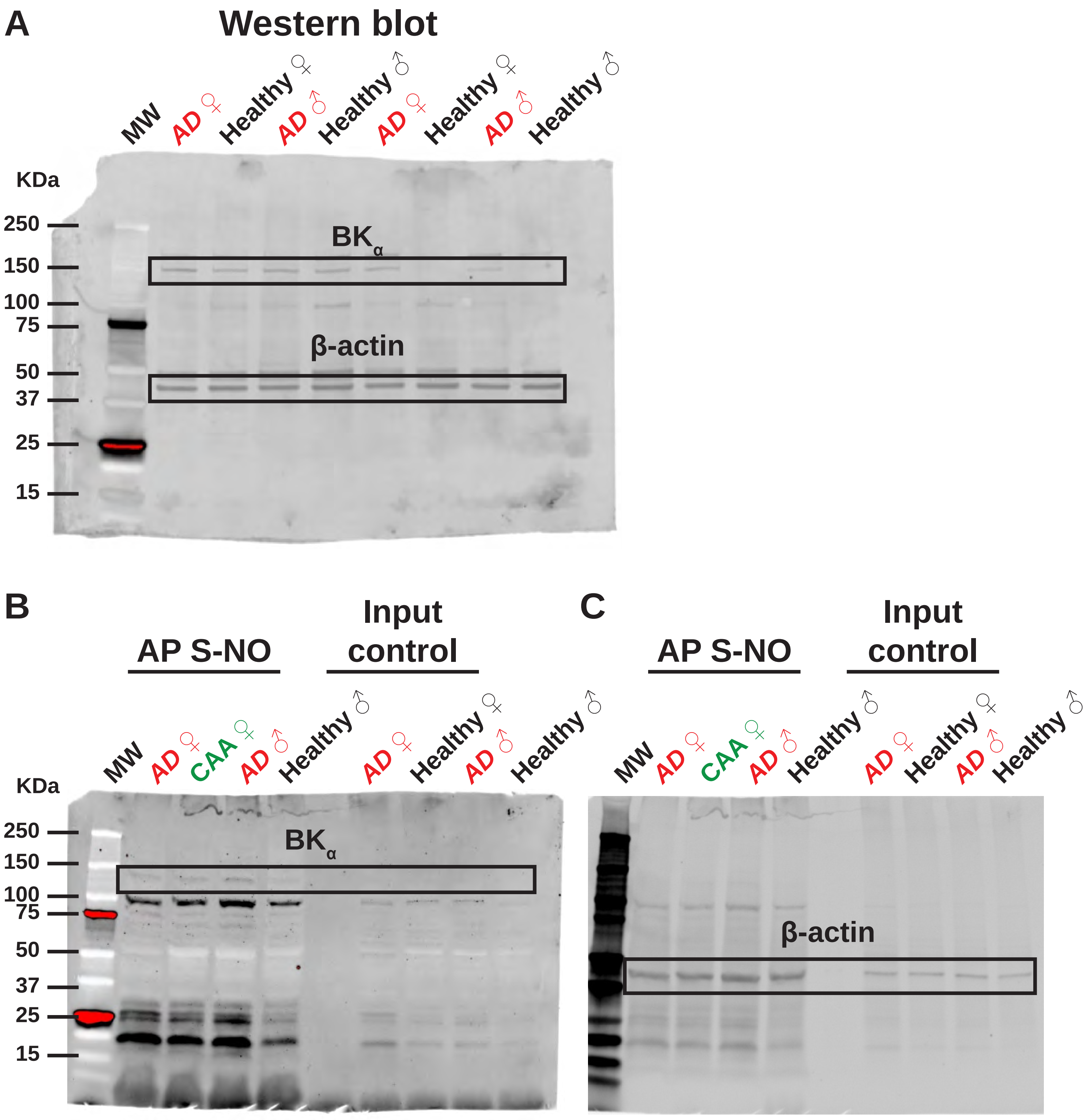

**Figure S12.** *Basal cortical perfusion assessed by laser speckle contrast imaging*

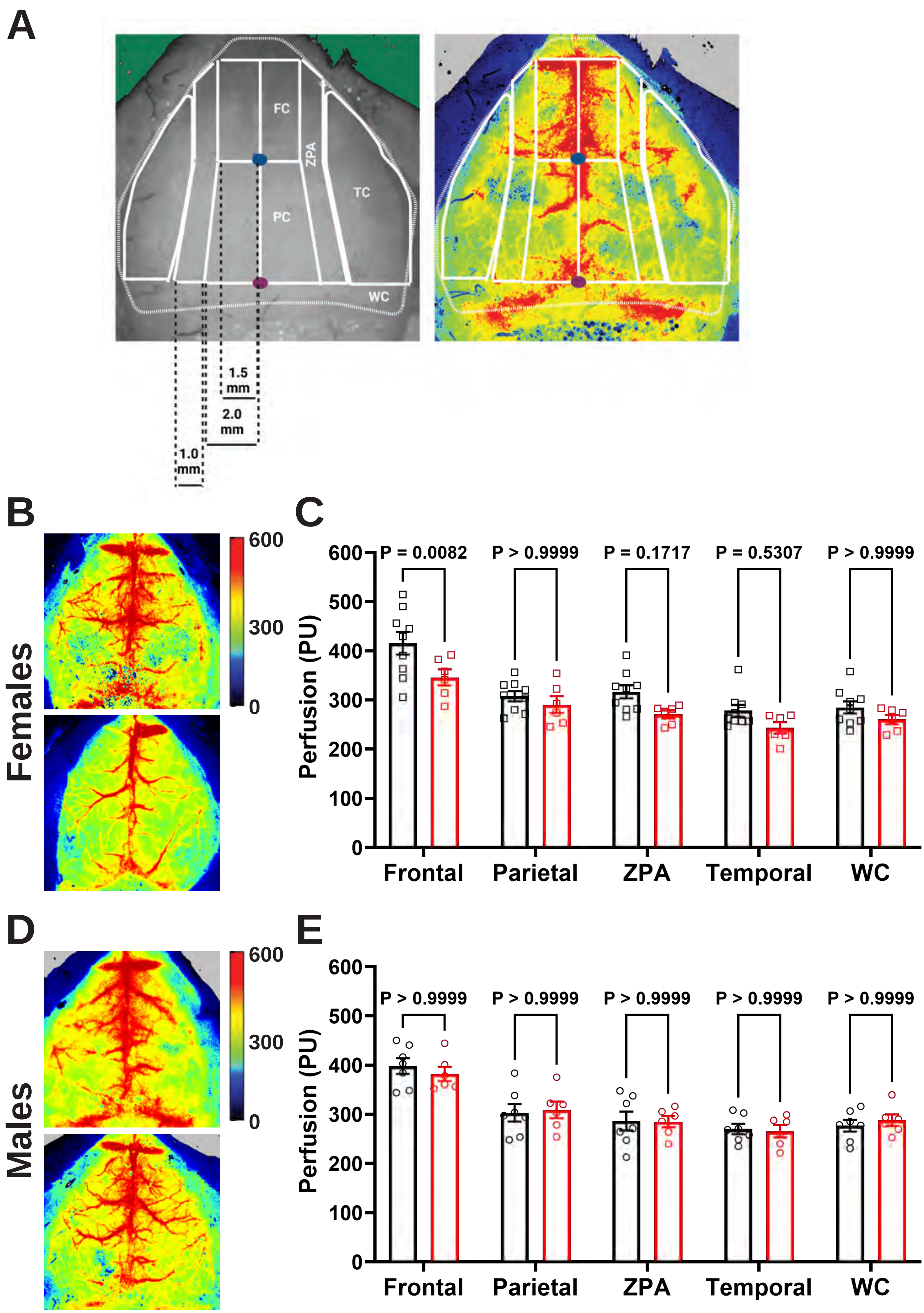

**Figure S13.** *Graphical summary of the main findings of the study*

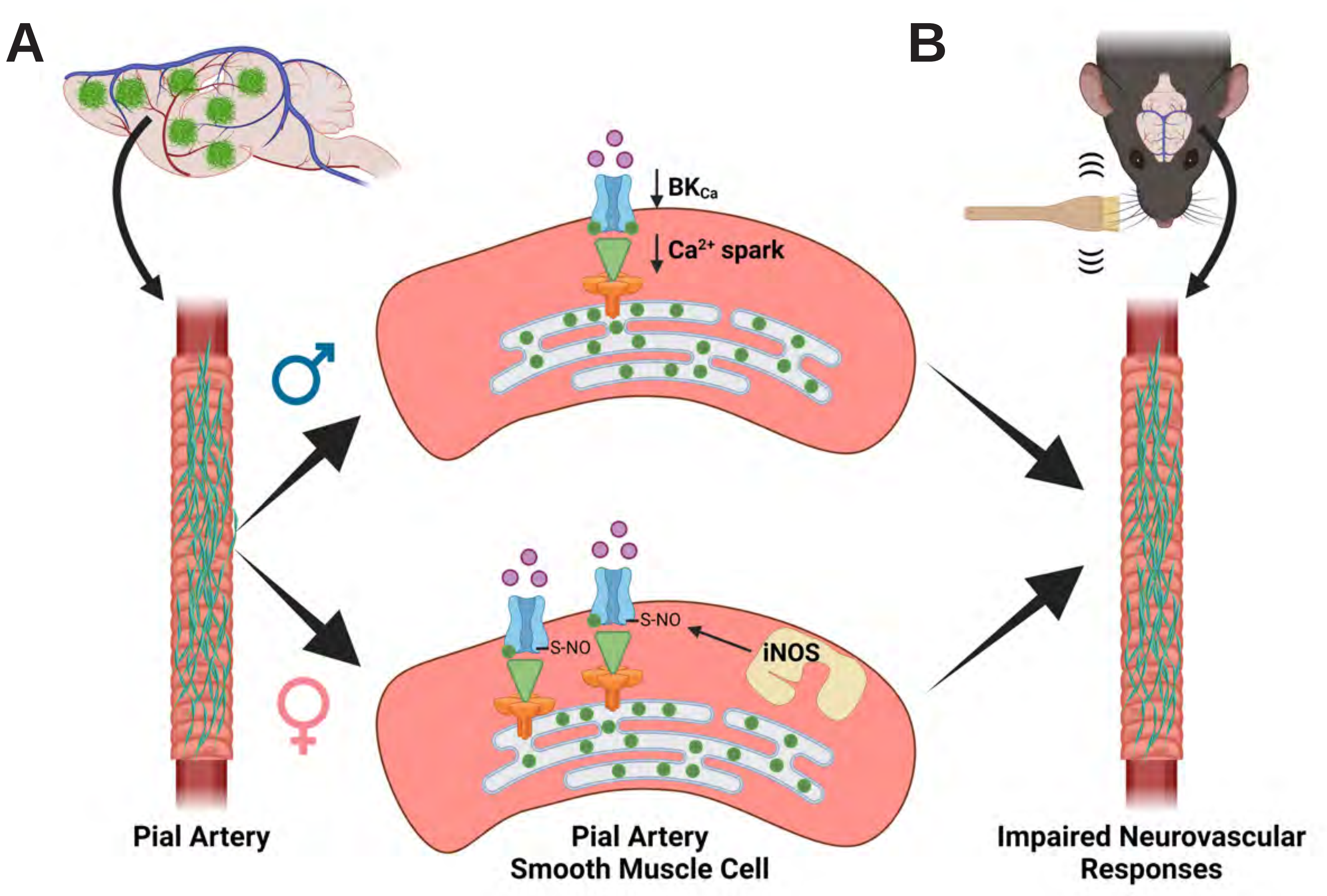
